# Interactions between the gut microbiome, dietary restriction, and aging in genetically diverse mice

**DOI:** 10.1101/2023.11.28.568137

**Authors:** Lev Litichevskiy, Maya Considine, Jasleen Gill, Vasuprada Shandar, Timothy O. Cox, Hélène C. Descamps, Kevin M. Wright, Kevin R. Amses, Lenka Dohnalová, Megan J. Liou, Monika Tetlak, Mario R. Galindo-Fiallos, Andrea C. Wong, Patrick Lundgren, Junwon Kim, Giulia T. Uhr, Ryan J. Rahman, Sydney Mason, Carter Merenstein, Frederic D. Bushman, Anil Raj, Fiona Harding, Zhenghao Chen, G.V. Prateek, Martin Mullis, Andrew G. Deighan, Laura Robinson, Ceylan Tanes, Kyle Bittinger, Meenakshi Chakraborty, Ami S. Bhatt, Hongzhe Li, Ian Barnett, Emily R. Davenport, Karl W. Broman, Robert L. Cohen, David Botstein, Adam Freund, Andrea Di Francesco, Gary A. Churchill, Mingyao Li, Christoph A. Thaiss

## Abstract

The intestinal microbiome changes with age, but the causes and consequences of microbiome aging remain unclear. Furthermore, the gut microbiome has been proposed to mediate the benefit of lifespan- extending interventions such as dietary restriction, but this hypothesis warrants further exploration. Here, by analyzing 2997 metagenomes collected longitudinally from 913 deeply phenotyped, genetically diverse mice, we provide new insights into the interplay between the microbiome, aging, dietary restriction, host genetics, and a wide range of health parameters. First, we find that microbiome uniqueness increases with age across datasets and species. Moreover, age-associated changes are better explained by cumulative exposure to stochastic events (neutral theory) than by the influence of an aging host (selection theory). Second, we unexpectedly find that the majority of microbiome features are significantly heritable and that the amount of variation explained by host genetics is as large as that of aging and dietary restriction. Third, we find that the intensity of dietary restriction parallels the extent of microbiome changes and that dietary restriction does not rejuvenate the microbiome. Lastly, we find that the microbiome is significantly associated with multiple health parameters — including body composition, immune parameters, and frailty — but not with lifespan. In summary, this large and multifaceted study sheds light on the factors influencing the microbiome and aspects of host physiology modulated by the microbiome.

## Introduction

Dietary restriction (DR) improves health and extends lifespan in diverse organisms^1,2^. However, its efficacy is highly variable and known to be influenced by numerous factors, including the type of dietary restriction and the organism’s genetic background^3–7^. To determine whether different types of dietary restriction extend lifespan in a genetically heterogeneous population such as humans, we randomized 960 Diversity Outbred mice to fasting and caloric restriction regimes and tracked their health and lifespan with extensive phenotyping. The design of this Dietary Restriction in Diversity Outbred mice (DRiDO) study is described in the parallel manuscript by Di Francesco et al.

A major goal of the DRiDO study was to identify predictors of longevity. One candidate for such a predictor is the gastrointestinal microbiome, which has recently been suggested to modulate aging^8–10^ as well as responses to dietary restriction^11–14^. To investigate the relationship between the gut microbiome and lifespan, we performed shotgun metagenomic sequencing on longitudinally collected stool samples. We generated 2997 metagenomic profiles from 913 Diversity Outbred mice, resulting in the largest-to-date mouse microbiome dataset. Using this dataset, we were able to address several fundamental questions.

First, how does the microbiome age? Numerous studies in mice^15–17^ and humans^18–21^ have reported age-associated microbiome changes, but these changes are inconsistent across cohorts^22^. Two community features frequently reported to increase with age are ɑ-diversity^18,22–24^ and uniqueness^25,26^. Whether these are universal properties of an aging microbiome remains unknown. Furthermore, it remains unclear to what degree age-associated microbiome changes are caused by the aging host.

Second, to what extent does host genetics influence the microbiome? The prevailing notion is that the environment, especially diet, has a much greater contribution to the gut microbiome than host genetics^27,28^. At the same time, multiple human studies have identified significant microbiome heritability and quantitative trait loci (QTLs) for microbiome features^29–33^. Moreover, a recent study in baboons found that nearly all microbiome features were significantly heritable but that identifying this heritability required large sample sizes and the use of longitudinal data^34^. Therefore, it remains unclear whether the influence of genetics on the microbiome is perhaps larger than previously appreciated.

Third, what aspects of host aging are modulated by the microbiome? Mice in the DRiDO study were deeply phenotyped, allowing us to ask whether the microbiome influences aspects of host physiology over the lifespan. The relationship between the microbiome and body composition is well-established^35–37^, but the DRiDO study was uniquely suited to discovering other host phenotypes influenced by the microbiome.

In addition to generating the metagenomic dataset in DO mice, we generated a second longitudinal microbiome dataset in genetically homogenous mice, performed a validation experiment to investigate the mechanism by which host age influences the microbiome, integrated our dataset with thousands of human metagenomic samples, and analyzed hundreds of longitudinal host phenotypes collected as part of the DRiDO study. We begin by describing the generation of the DRiDO microbiome dataset.

### Longitudinal metagenomic sequencing of Diversity Outbred mice

The design of the DRiDO study is described in depth in Di Francesco et al. Briefly, 937 female DO mice began one of five dietary interventions at six months of age (**Fig. 1a**, see Methods): *ad libitum* food consumption (AL, control group), fasting one day per week (1D), fasting two consecutive days per week (2D), consuming 20% fewer calories every day (20), or consuming 40% fewer calories every day (40). Mice were extensively phenotyped over their lifespans (**Extended Data Table 1**). All dietary restriction interventions significantly extended lifespan (log-rank test, p < 2.2e-16), but there was substantial inter-individual variability (**Fig. 1b**). The largest extension in lifespan was achieved by the 40% caloric restriction group (36% increase in median lifespan versus AL).

**Fig. 1.**
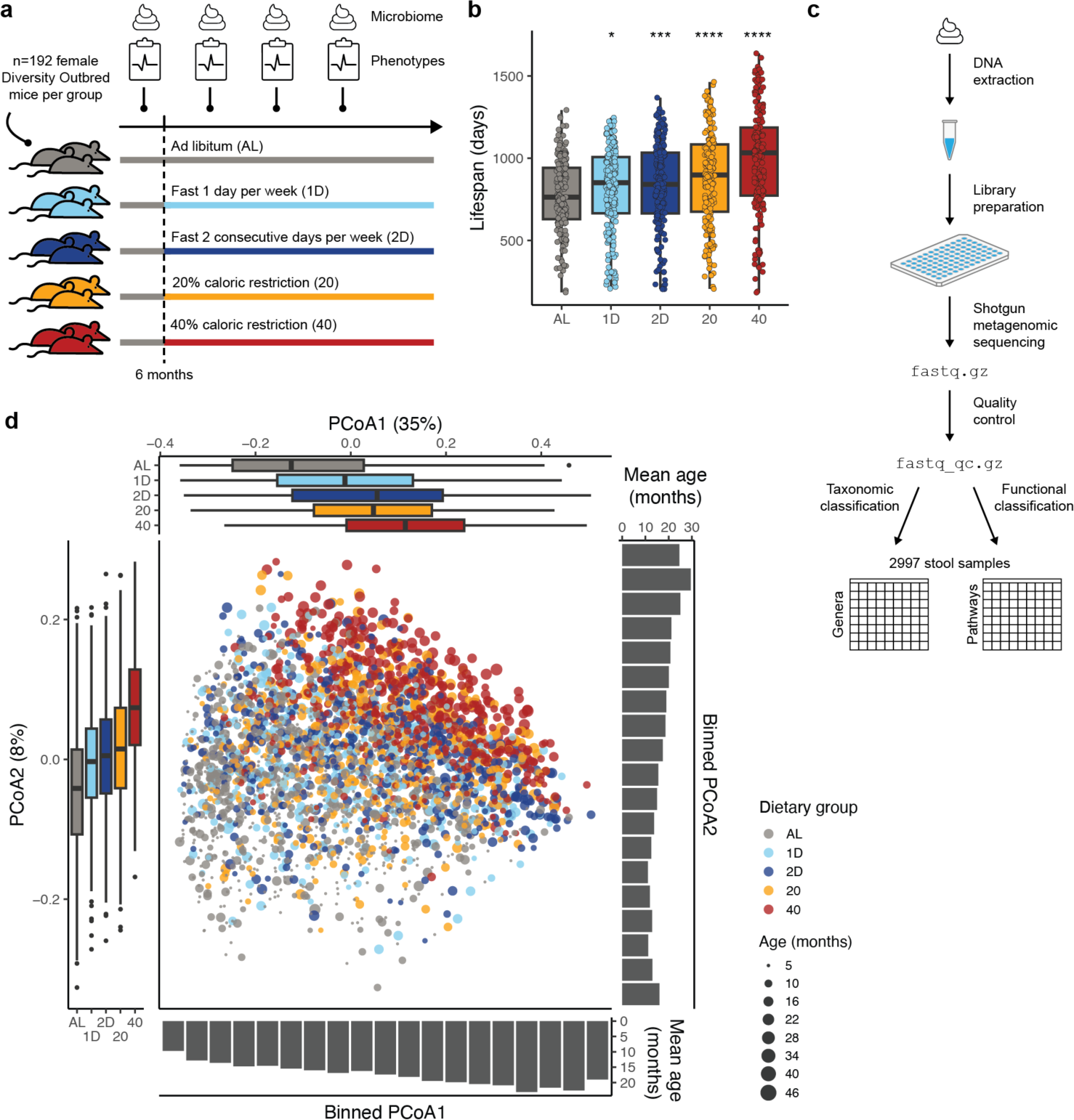
Overview of DRiDO study and microbiome dataset. a, At six months of age, genetically diverse mice started one of five dietary interventions. They were extensively phenotyped and stool was collected for microbiome profiling. **b**, Lifespan per dietary group. Each dot is one of 924 mice: 23 mice died before the start of dietary restriction and 13 mice died from accidental mishandling. p-values were calculated with pairwise log-rank tests against the AL group. **c**, Microbiome data generation consisted of extracting DNA from stool samples, preparing libraries, performing shotgun metagenomic sequencing, performing quality control, and finally taxonomic and functional classification. After all quality control, the cohort consisted of 2997 stool samples. **d**, Principal Coordinates Analysis (PCoA) plot of quality-controlled microbiome samples. Ordination based on Bray-Curtis distances of genus-level relative abundances. Color denotes dietary group, and size encodes mouse age at the time of stool collection. Boxplots along the sides show PCoA1 (top) and PCoA2 (left) coordinates per dietary group. Barplots along the sides show the mean age of stool samples within each bin of PCoA1 (bottom) and PCoA2 (right) coordinates. PCoA1 and PCoA2 explain 35% and 8% of overall variance, respectively.

To characterize the gut microbiome, we collected stool samples approximately every six months and performed paired-end (2x150 bp) shotgun metagenomic sequencing on extracted DNA (**Fig. 1c**). We sequenced 3586 stool samples (mean 14.1M read pairs per sample), 62 positive controls, and 71 negative controls (**Extended Data Fig. 1**a-c). Different library preparations of the same DNA and repeat sequencing of the same libraries produced highly similar microbiome profiles (**Extended Data Fig. 1**d, e).

In addition, we leveraged^38^ the fact that every mouse in our study was genetically unique and that each stool sample contained some host DNA (∼9% of reads) to exclude samples where the genotype of the host and stool sample did not definitively match (**Extended Data Fig. 2**a-d, **Supplementary Note 1, Supplementary Tables 1-2**). Our pipeline for identifying sample mix-ups allowed us to detect and remedy errors that occurred during data generation (**Extended Data Fig. 2**e), including an animal swap (**Extended Data Fig. 2**f, g).

Samples were also discarded if they had inconsistent metadata, insufficient reads, an unusually high proportion of host reads, or if they appeared to be outliers (**Extended Data Fig. 3**a, see Methods). Our final quality-controlled cohort consisted of 2997 metagenomic profiles from 913 mice, with a median of three samples per mouse, five timepoints with a large number (at least 360) of samples, and at least 550 samples per dietary group.

We performed taxonomic classification using Kraken2 (ref. ^39^) and the Mouse Gastrointestinal Bacterial Catalogue^40^ (MGBC) as reference, and we performed functional classification with HUMAnN3 (ref. ^41^). We used Kraken2 and MGBC instead of the taxonomic results available from HUMAnN3 for two reasons: 1) Kraken2 with MGBC classified more reads (**Extended Data Fig. 3**b) and 2) the fraction of characterized (i.e., named) taxa was higher (55% versus 14% for genera, 12% versus 9% for species). The two methods showed good concordance (**Extended Data Fig. 3**c). We hereafter present Kraken2 taxonomic results, except in the comparisons to other datasets. For functional results, we present MetaCyc^42^ pathways and further distinguish between “community-wide” and “specialized” pathways, in which specialized pathways are defined as being highly correlated with just a few genera (**Extended Data Fig. 3**d).

### Most microbiome features change with age

First, we assessed whether aging has an effect on the microbiome. Two-dimensional ordination plots of both genera (**Fig. 1d**) and pathways (**Extended Data Fig. 3**e) suggested a strong effect of host age influencing overall microbiota composition and function (PERMANOVA pseudo-F = 67.0, p-value = 0.001, df = 1 for genera; pseudo-F = 44.7, p-value = 0.001, df = 1 for pathways). To assess the impact of aging on individual microbiome features, we fit a linear mixed model separately for each feature (see Methods). This model accounted for age, dietary group, host genetics (via a kinship matrix), mouse identity, and technical factors including DO mouse breeding cohort, the cage in which a mouse was housed, and DNA extraction batch.

Most genera and pathways were significantly (conditional Wald test, adjusted p-value < 0.01) associated with host age (**Fig. 2a**). Genera whose relative abundance increased most strongly with age included *Bifidobacterium* (**Fig. 2b**), *Turicibacter*, and *Alistipes*, while genera with the greatest decreases included poorly characterized microbes such as ASF356, UMGS268, and UBA9475. The community feature with the strongest positive association with host age was uniqueness – defined as a microbiome sample’s β-diversity (or distance) to its nearest neighbor^26^ – based on both genera (**Fig. 2c**) and pathways (**Extended Data Fig. 4**a). We confirmed that this trend was present in all dietary groups (**Extended Data Fig. 4**b) and that it was not due to the number of mice per cage decreasing with age (**Extended Data Fig. 4**c). We found that ɑ-diversity (as measured by Shannon and Simpson indexes) increased with age, but the trend was not significant (**Extended Data Fig. 4**d).

**Fig. 2.**
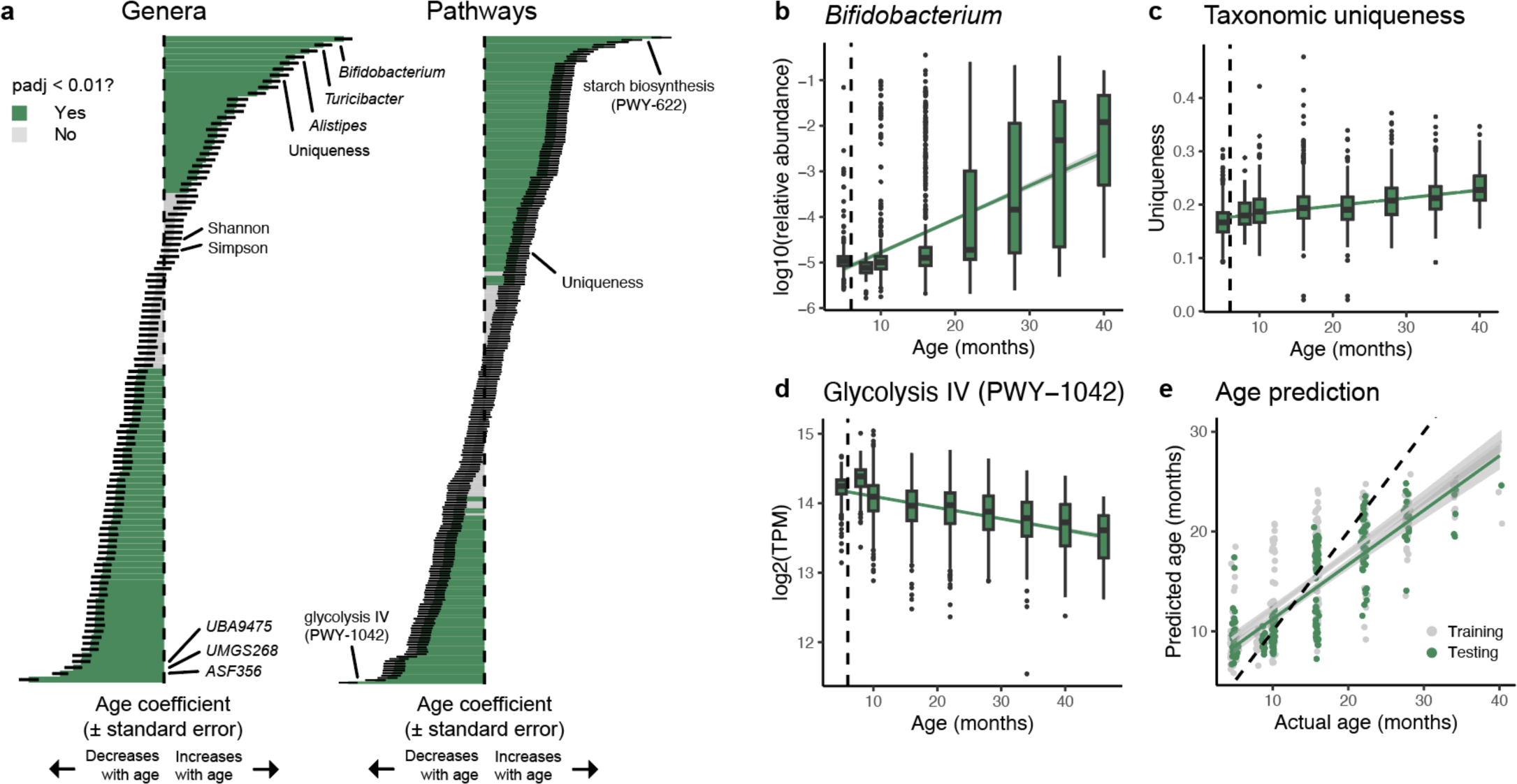
**Age-associated microbiome changes in DO mice. a**, Effect of age on genera (left) and pathways (right). Age coefficients were calculated with a linear mixed model. The thin black lines show standard error around the mean. Green indicates an adjusted p-value < 0.01. **b,** *Bifidobacterium* increases with age. Green line represents line of best fit and 95% confidence interval. Vertical dashed line at six months represents start of dietary restriction. **c**, Taxonomic uniqueness increases with age. Uniqueness is defined as the minimum distance to any other microbiome sample. **d**, Glycolysis IV (PWY-1042) decreases with age. **e**, Host age prediction using microbiome profiles. This analysis considered genus-level relative abundance profiles from AL mice. Gray dots are out-of-bag predictions on training data (70%). Green dots are predictions by the classifier on never-before-seen testing data (30%). Gray and green lines represent lines of best fit and 95% confidence intervals for training and testing data, respectively. Black dotted line at y=x represents perfect accuracy.

Many microbial pathways were affected by host age. The pathway with the largest positive association with age was starch biosynthesis (PWY-622), a specialized pathway that was strongly correlated with *Bifidobacterium* (**Extended Data Fig. 4**e). The pathway with the largest negative association was glycolysis IV (PWY-1042, **Fig. 2d**), a community-wide pathway.

Since many microbiome features were associated with host age, we asked whether microbiome information could be used to predict host age^16,21,43–45^ (see Methods). Considering just AL mice, we found that a random forest classifier could predict host age using either genera (**Fig. 2e**) or pathways (**Extended Data Fig. 4**f), demonstrating that the gut microbiome undergoes age-associated changes that can be detected by a machine-learning algorithm. However, the mean absolute error on held-out samples was high (15.7 ± 13.4 weeks and 22.4 ± 16.0 weeks for genera and pathways, respectively), indicating that additional factors influence microbiome composition and function.

### Universality of age-associated microbiome changes

To investigate whether any of these age-associated changes constituted a conserved microbiome signature of aging, we compared our dataset to other aging microbiome datasets. We compared AL mice from the DO cohort to a longitudinal mouse microbiome study that we conducted in male C57BL/6 mice and to a publicly available human metagenomic sequencing database^46^ (**Fig. 3a**). Because the human samples had been processed with HUMAnN, we processed the C57BL/6 dataset with HUMAnN and used our taxonomic HUMAnN results for DO mice to enable more direct comparisons. We fit linear mixed models separately for each dataset to identify age-associated taxonomic and functional features (see Methods).

**Fig. 3.**
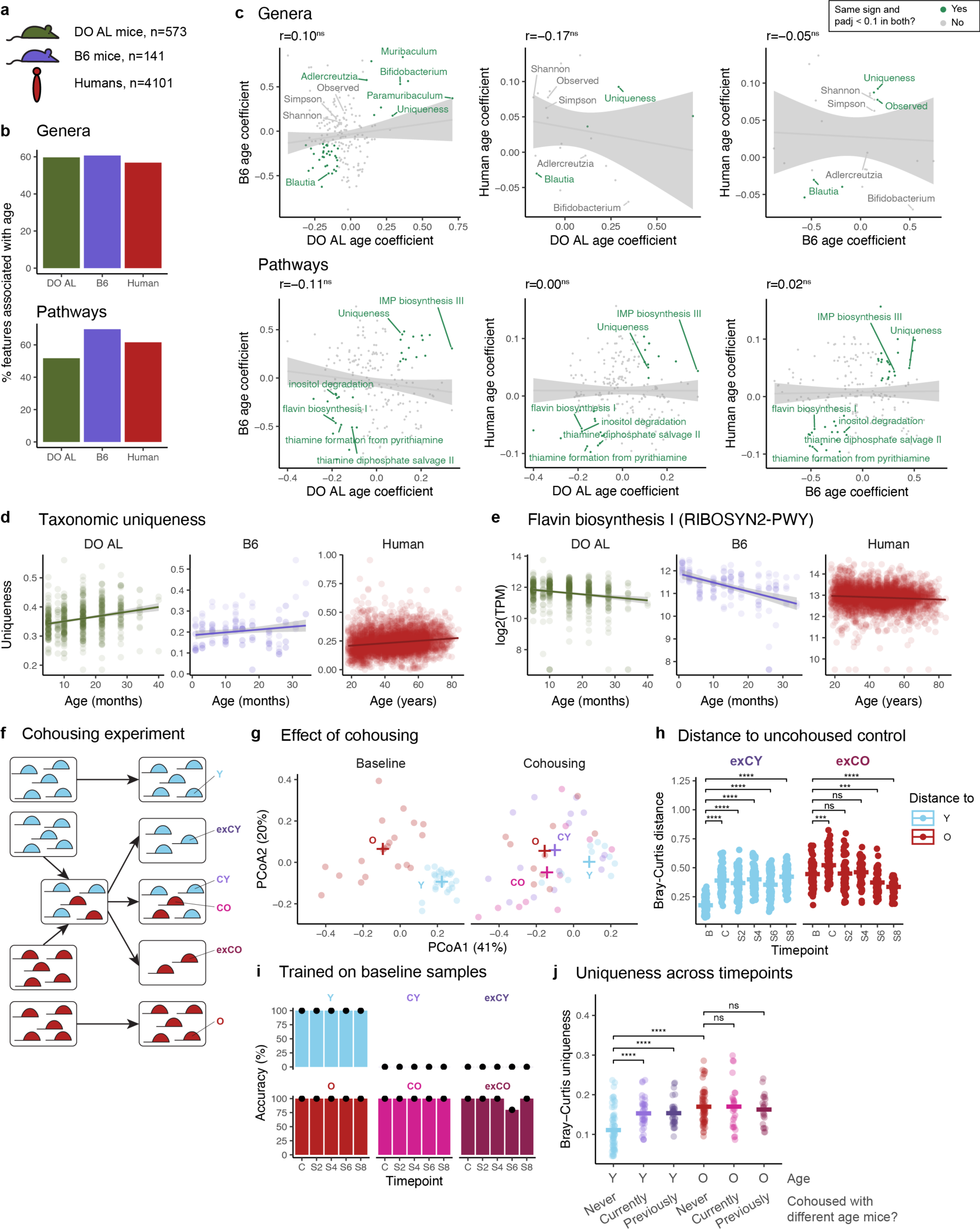
**Universality of age-associated microbiome changes. a**, We compared 573 samples from DO AL mice to 141 samples from a different mouse aging cohort (“B6”) to 4101 human gut microbiome samples. **b**, Percentage of features significantly associated with age (adjusted p-value < 0.1) within each dataset for genera (top) and pathways (bottom). **c**, Comparison of age-associated taxonomic (top) and functional (bottom) changes across datasets. Each pairwise comparison shows all features that passed prevalence filtration in both datasets. Line of best fit and 95% confidence interval shown in gray. Pearson correlation and corresponding p-value shown above each plot. Features significantly associated with age and with the same sign in the pairwise comparison are shown in green. **d**, Taxonomic uniqueness increases with age in all three datasets. **e**, Flavin biosynthesis I (RIBOSYN2-PWY) decreases with age in all three datasets. **f**, Schematic of cohousing and separation experiment. Young mice are blue, old mice are red. Y = young always housed with young, O = old always housed with old, CY = young housed with old, CO = old housed with young, exCY = formerly CY that were separated from old, and exCO = formerly CO that were separated from young. **g**, PCoA of samples at baseline and after one month of cohousing. Ordination based on all samples shown in this plot. + denotes group centroid. **h**, Bray-Curtis distances between previously cohoused mice (exCY, exCO) and non-cohoused controls (Y, O). Significance of group differences was evaluated with a t-test. **i**, Random forest classifier trained on baseline samples and evaluated on cohousing and separation samples. Accuracy is the percentage of samples within each group correctly classified as young or old. **j**, Uniqueness split by age and cohousing status. Significance of group differences was evaluated with a t-test. B = baseline, C = cohousing, S2 = 2 weeks of separation, S4 = 4 weeks of separation, etc.

Within each dataset, many genera and pathways were significantly associated with age (**Fig. 3b**), but among taxonomic features that could be compared across datasets, there was little concordance in age-associated changes (**Fig. 3c**). Just one taxonomic feature significantly increased with age in all three datasets: uniqueness (**Fig. 3d**). We verified that uniqueness increased with age in most of the studies comprising our human cohort (**Extended Data Fig. 5**a). *Blautia* was negatively associated with age across all datasets, but upon closer inspection, the association in humans was inconsistent (**Extended Data Fig. 5**b). Some genera — such as *Paramuribaculum*, *Muribaculum*, and *Adlercreutzia* — significantly increased with age in both mouse cohorts but not in humans. No ɑ-diversity metrics were significantly associated with age in all three datasets and, in humans, the relationship between ɑ-diversity and age was highly variable by study (**Extended Data Fig. 5**c). Taken together, the only taxonomic signature of aging that we could identify was an increase in uniqueness.

As with genera, there was poor overall concordance of age-associated pathway changes across datasets. Nevertheless, several pathways showed consistent changes: inosine-5’-phosphate (IMP) biosynthesis (PWY- 7234) increased with age, while thiamine diphosphate formation (PWY-7357), thiamine diphosphate salvage (PWY-6897), inositol degradation (PWY-7237), and flavin biosynthesis (RIBOSYN2-PWY, **Fig. 3e**) decreased with age. With the exception of IMP biosynthesis, these are all community-wide pathways (**Extended Data Fig. 5**d). In addition, functional uniqueness increased with age in all three datasets. These results suggest that there may exist a functional signature of microbiome aging characterized by decreased production of cofactors such as thiamine (vitamin B1) and riboflavin (vitamin B2) and increased production of IMP, a necessary precursor for de novo purine biosynthesis.

### Microbiome changes over the lifespan reflect cumulative stochastic exposure, not host age

Thus far, we had identified a strong but inconsistent effect of age on the microbiome. This motivated us to ask what causes the microbiome to change with age. We investigated this question through the lens of selection and neutral theory^47–49^. We hypothesized that the apparent “age” of the microbiome is dictated by (a) selective pressure based on host age (selection theory), or (b) exposure to stochastic events that accumulate with time (neutral theory). To test these hypotheses, we needed to disentangle host age from microbiome age. We did this by cohousing young (8 weeks) and old (19-20 months) C57BL/6 mice for one month and monitoring their microbiomes (with 16S sequencing) after 2, 4, 6, and 8 weeks of separation (**Fig. 3f**). Cohousing caused the microbiomes of young mice to resemble those of old mice (**Fig. 3g**). In other words, it had decoupled host age and microbiome age in young mice. After separation, the microbiomes of young mice never reverted to a young state: they remained significantly different from the microbiomes of non-cohoused young mice (**Fig. 3h**). Furthermore, a classifier trained on baseline microbiome samples always incorrectly identified both cohoused and previously-cohoused young mice as old (**Fig. 3i**). These results indicate that a young host does not dictate the apparent age of the microbiome. The effect of cohousing was much less pronounced in old mice (**Fig. 3g, h**), and the classifier always correctly identified old mice, even during cohousing (**Fig. 3i**).

We observed again that uniqueness was higher in old compared to young mice (**Fig. 3j**). Notably, young mice cohoused with old mice acquired higher microbiome uniqueness compared to non-cohoused young mice, and this was not reversed upon separation from old mice. Thus, uniqueness appears to reflect the cumulative stochastic exposure of the microbial community, rather than host age. More broadly, these findings suggest that microbiome aging is better explained by neutral theory, with age-associated microbiome changes resulting from within-community development over time, rather than from host influence.

### Host genetics shape the gut microbiome

The DRiDO study’s use of genetically diverse and genotyped mice enabled us to evaluate the contribution of host genetics to microbiome composition and function. The first indication that host genetics might have a substantial influence on the microbiome was the observation that (genetically diverse) DO mice had higher uniqueness than (genetically homogeneous) B6 mice (**Fig. 3d**; t-test, t = 23.226, df = 174.16, p-value < 2.2e- 16), suggesting that genetic diversity led to more inter-individual microbiome variation.

We found that the majority of genera (66%, mean heritability of heritable features = 0.11) and pathways (51%, mean = 0.08) had significant, though modest, heritability (**Fig. 4a**, see Methods). The most heritable taxa were *Lactobacillus*, *Parvibacter,* and a novel genus in the Erysipelatoclostridiaceae family. The most heritable microbial pathway was lactose and galactose degradation (LACTOSECAT-PWY), a specialized pathway strongly correlated with *Lactobacillus* (**Extended Data Fig. 6**a). Heritable community-wide pathways include de novo queuosine biosynthesis (PWY-6700) and fatty acid biosynthesis initiation (PWY66-429).

**Fig. 4.**
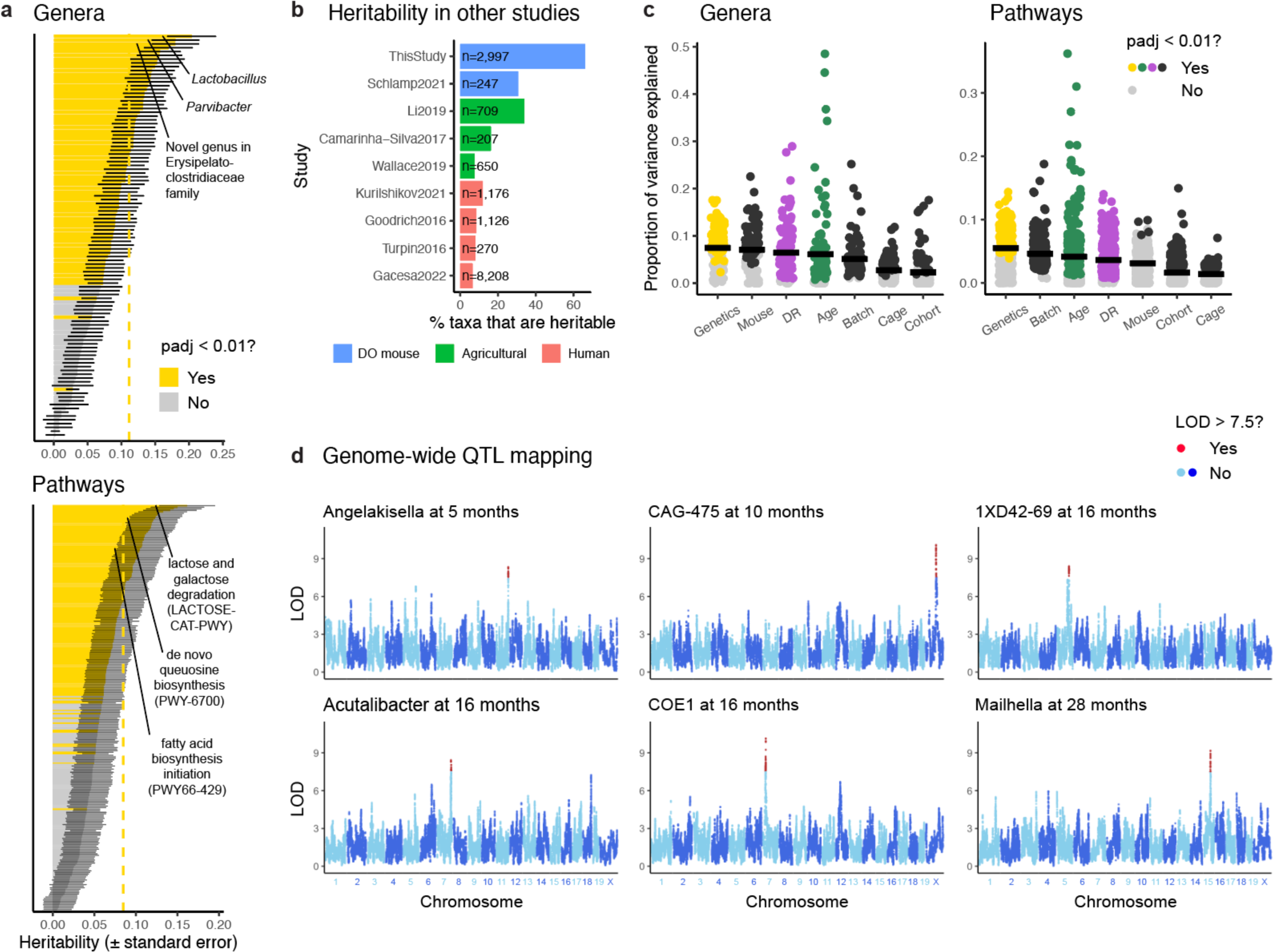
**Genetic influence on the microbiome. a**, Heritability of genera (top) and pathways (bottom). For visual clarity, we omitted 9 pathways with heritability and standard error approximately equal to zero. Vertical dashed line shows mean heritability for heritable features. **b**, Percentage of taxa with significant heritability (as reported by the authors) in other studies. The number of samples per study is indicated. The color of each bar indicates whether the study was performed in Diversity Outbred mice, agricultural animals, or humans. **c**, Proportion of variance explained (PVE) by various experimental variables for genera (left) and pathways (right). Horizontal lines show the mean PVE. **d**, Genome-wide results for the six age-specific significant QTLs with adjusted p-value less than 0.1. Markers with LOD greater than 7.5 are colored red.

Since human studies often report a minor role of genetics in shaping the gut microbiome^27,31^, we confirmed our surprising result with several orthogonal approaches. First, we made sure that this was not a peculiarity of the software we used for calculating heritability by exactly reproducing our heritability estimates with an alternative software package (**Extended Data Fig. 6**b). Second, we compared our results to an independent dataset of DO mice^50^ and found that, of genera that could be compared across studies, the most heritable genus in both datasets was *Lactobacillus* (**Extended Data Fig. 6**c, see Methods). Third, we calculated heritability separately per age (see Methods) to determine whether our widespread heritability was a consequence of using longitudinal data, as previously suggested^34^. Many more features had significant heritability when using longitudinal data compared to cross-sectional data, but downsampling to a similar number of samples as in cross-sectional data completely erased heritability (**Extended Data Fig. 6**d). In other words, a large number of samples was critical for detecting heritability, not the use of longitudinal data *per se*.

Another potential explanation for why we observed higher heritability than typically reported in human studies relates to differences in environmental variability. Heritability is a relative measure; it is defined as VG / (VG + VE), where VG is the amount of variance explained by additive genetic effects, and VE is all other sources of variance. Therefore, larger VE will decrease heritability even if VG is unchanged. If we assume that human observational studies have higher VE (i.e., more unexplained environmental variability) than studies involving agricultural animals and even higher VE than studies in laboratory animals, then differences in VE may partially explain why heritability estimates are generally lower in humans than in agricultural animals and lower still than in DO mice (**Fig. 4b**).

How does the magnitude of this genetic effect compare to the effects of other experimental variables? To answer this question, we fit a linear mixed model in which all variables were treated as random effects (see Methods). We found that the proportion of variance explained by genetics was similar to that of dietary restriction and aging (**Fig. 4c**). The majority of variance remained unexplained (63% for genera, 76% for pathways), emphasizing that the microbiome is strongly influenced by factors that are yet unaccounted for. Technical factors such as cage, cohort of DO breeding, and DNA extraction batch explained smaller, but still significant, amounts of variance, emphasizing the importance of retaining these technical factors in our linear modelling.

Lastly, because many microbiome features showed significant heritability, we performed genome-wide quantitative trait loci (QTL) mapping to find loci that influence microbiome composition (see Methods). We tested 107 features at five different ages and identified just six significant QTLs (**Fig. 4d**, **Extended Data Table 2**). The disconnect between widespread microbiome heritability but few significant QTLs likely reflects the fact that microbial abundance is a complex, polygenic trait. Furthermore, these six QTLs were significant at only one age (**Extended Data Fig. 6**e), suggesting that the genetic influence on the microbiome may be temporally variable.

### Effects of dietary restriction

In addition to host age and genetics, dietary restriction also had a strong influence on the microbiome (**Fig. 1d**, **Fig. 4c**). Indeed, we found that nearly all microbiome features were affected by DR (**Fig. 5a**). We recovered the commonly observed phenomenon^17,51–53^ of DR increasing the abundance of *Lactobacillus* and closely related genera (**Fig. 5b**). We also found that ɑ-diversity is increased by DR (except 2D fasting, **Fig. 5c**), and uniqueness is increased by DR (**Fig. 5d**), in addition to increasing with age. The microbial functions most affected by DR were specialized pathways (**Extended Data Fig. 7**a), such as lysine biosynthesis (PWY-2941, highly correlated with *Ligilactobacillus*) and O-antigen building blocks biosynthesis (OANTIGEN-PWY, highly correlated with *Lactobacillus*). Community-wide pathways affected by DR included the urea cycle (PWY-4984, **Fig. 5e**) and citrulline biosynthesis (**Extended Data Fig. 7**b).

**Fig. 5.**
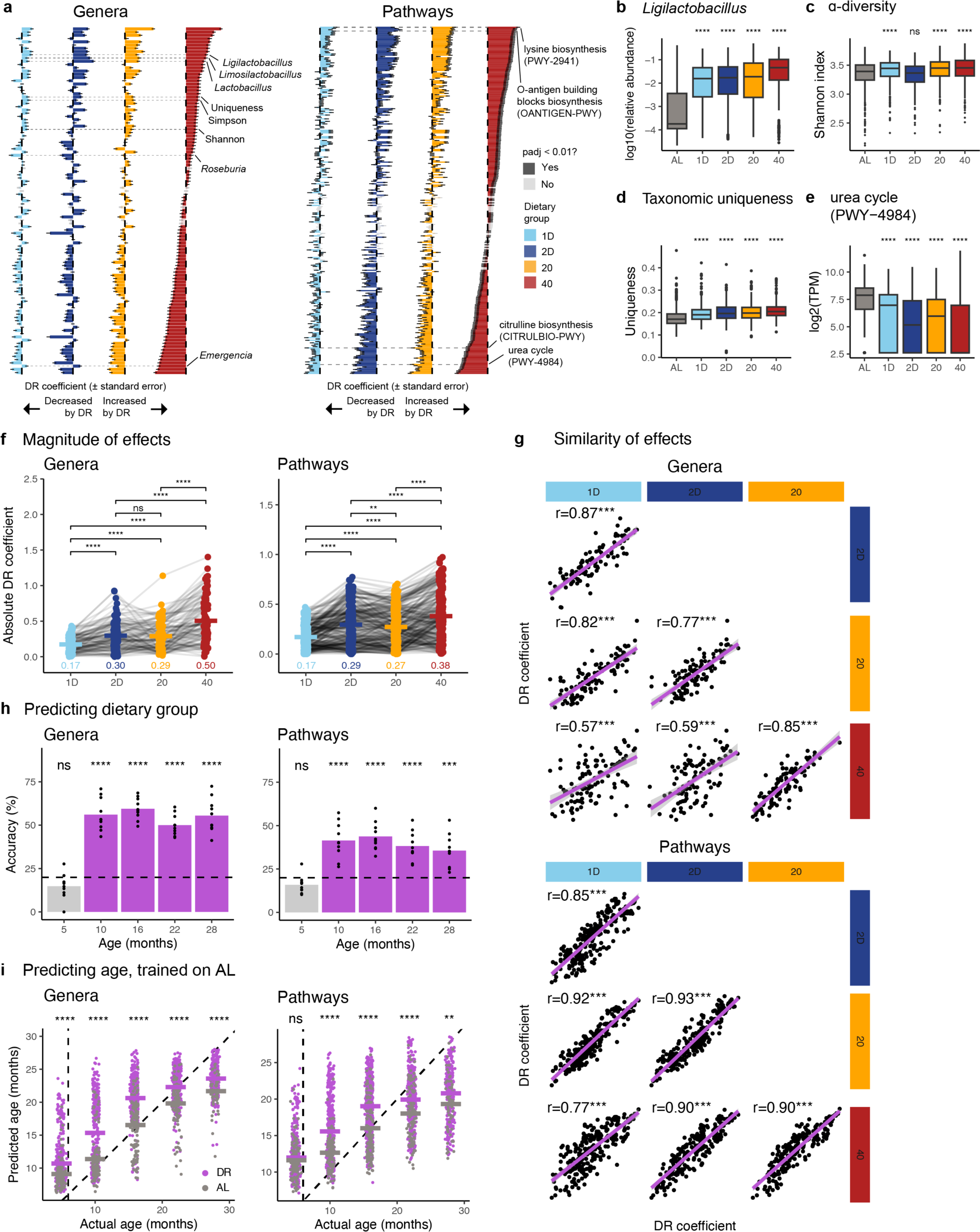
**Effects of dietary restriction on the microbiome. a**, Effect of dietary restriction (DR) on genera (left) and pathways (right). Horizontal dashed gray lines are visual aids to help compare across dietary groups. **b**, *Ligilactobacillus* was increased by DR. Statistical significance evaluated by a t-test against the AL group. **c**, ɑ-diversity (as measured by Shannon index) was increased by all DR groups except 2D fasting. **d**, Taxonomic uniqueness was increased by DR. **e**, The urea cycle (PWY-4984) was decreased by DR. **f**, Absolute magnitude of DR coefficients for genera (left) and pathways (right). Each dot is a microbiome feature, gray lines connect the same feature. Horizontal bars show the mean. Statistical significance evaluated by a paired t-test. **g**, Comparison of DR coefficients. Each dot is a genus (top) or pathway (bottom). Pearson correlation and p-value is indicated above each scatterplot. Line of best fit and 95% confidence interval are shown in purple. **h**, Predicting dietary group with genera (left) or pathways (right). Each dot represents prediction accuracy in one cross-validation fold. Horizontal dashed line at 20% represents expected accuracy by chance. Statistical significance evaluated by a one-sided t-test (testing whether the mean accuracy is greater than 20%). **i**, Age prediction with a random forest classifier trained on AL samples. Input to classifier was either genera (left) or pathways (right). Vertical dashed line at six months represents start of dietary restriction, diagonal dashed line represents perfect prediction. Statistical significance evaluated by a t-test between AL and DR predictions at each age.

We observed that 40% CR had the strongest overall effect on the microbiome, followed by 2D fasting and 20% CR, and finally by 1D fasting, suggesting that the intensity of dietary restriction parallels the extent of microbiome changes (**Fig. 5f**). To determine whether fasting and CR have similar global effects on the microbiome, we correlated dietary restriction coefficients calculated by the linear mixed model (**Fig. 5g**). For both genera and pathways, dietary coefficients from different DR groups were highly correlated, but correlations were higher for pathways. For genera, the correlations within CR and fasting groups (i.e., 20% to 40%, 1D to 2D) were higher than the correlations across CR and fasting groups (i.e., 20% to 1D, 20% to 2D, 40% to 1D, 40% to 2D). This pattern was not observed for pathways. Two examples of taxonomic changes specific to CR or fasting are *Emergencia —* which was unaffected by fasting but decreased by CR — and *Roseburia* — which was decreased only by fasting (**Extended Data Fig. 7**c). We were unable to find pathways differentially affected by CR or fasting (**Extended Data Fig. 7**d). These findings suggest that, overall, CR and fasting have similar effects on microbial composition and function, but the effects on composition are more variable and more specific to the type of DR.

As an orthogonal way to investigate how the gut microbiome is influenced by dietary restriction, we asked whether a machine-learning algorithm would be able to predict the dietary group of a mouse based on its microbiome profile. We trained a random forest classifier (separately at each age) to predict dietary group (see Methods). As expected, the classifier performed no better than chance (20% accuracy) at five months (i.e., prior to the start of dietary restriction) for both genera and pathways (**Fig. 5h**). After the initiation of dietary restriction, the classifier performed significantly better than chance. Accuracy was higher for genera than for pathways (t-test between genus and pathway accuracies after five months, p=9.6e-11), consistent with the idea that CR and fasting induce more distinct taxonomic changes than functional changes. Furthermore, accuracy was highest for the AL, 2D, and 40% dietary groups (**Extended Data Fig. 7**e), also consistent with our finding that a more intense dietary intervention creates a more distinct microbiome state.

Because DR extended lifespan and improved various health parameters (see manuscript by Di Francesco et al.), we assessed whether DR induced a more youthful microbiome state. To answer this question, we trained a random forest classifier on all AL samples and predicted the host age of DR samples. If DR produced a more youthful microbiome state, the predicted age of DR samples would be lower than the predicted age of AL samples. Surprisingly, we found that DR samples had higher (t-test, p-value < 0.01) predicted ages than AL samples (**Fig. 5i**). Conversely, a classifier trained on all 40% CR samples predicted lower age for AL samples than for the other DR samples (**Extended Data Fig. 7**f). Furthermore, two-dimensional ordination showed that aging and DR shifted the microbiome in the same direction (**Extended Data Fig. 7**g). In summary, we find that more intense dietary interventions cause larger microbiome changes, that CR and fasting have more concordant functional effects than taxonomic effects, and that DR does not “rejuvenate” the microbiome to a more youthful state.

### The microbiome influences host physiology but not lifespan

Having characterized the factors that influence the microbiome, we next asked whether the microbiome modulates any of the host phenotypes measured in the DRiDO study. We tested genera and pathways for association with 197 phenotypic traits across 13 assays, while controlling for age, dietary group, and mouse (**Fig. 6a**, see Methods). The proportion of significant (adjusted p-value < 0.01) associations was similar for genera (0.9%) and pathways (0.7%, **Extended Data Fig. 8**). We observed associations with phenotypes measured in the body weight, body composition (dual-energy X-ray absorptiometry, DEXA), metabolic cage, frailty, and flow cytometry assays (**Fig. 6b**). No associations were observed with phenotypes measured in the echocardiogram, glucose, grip strength, void (bladder function), rotarod, or wheel running assays.

**Fig. 6.**
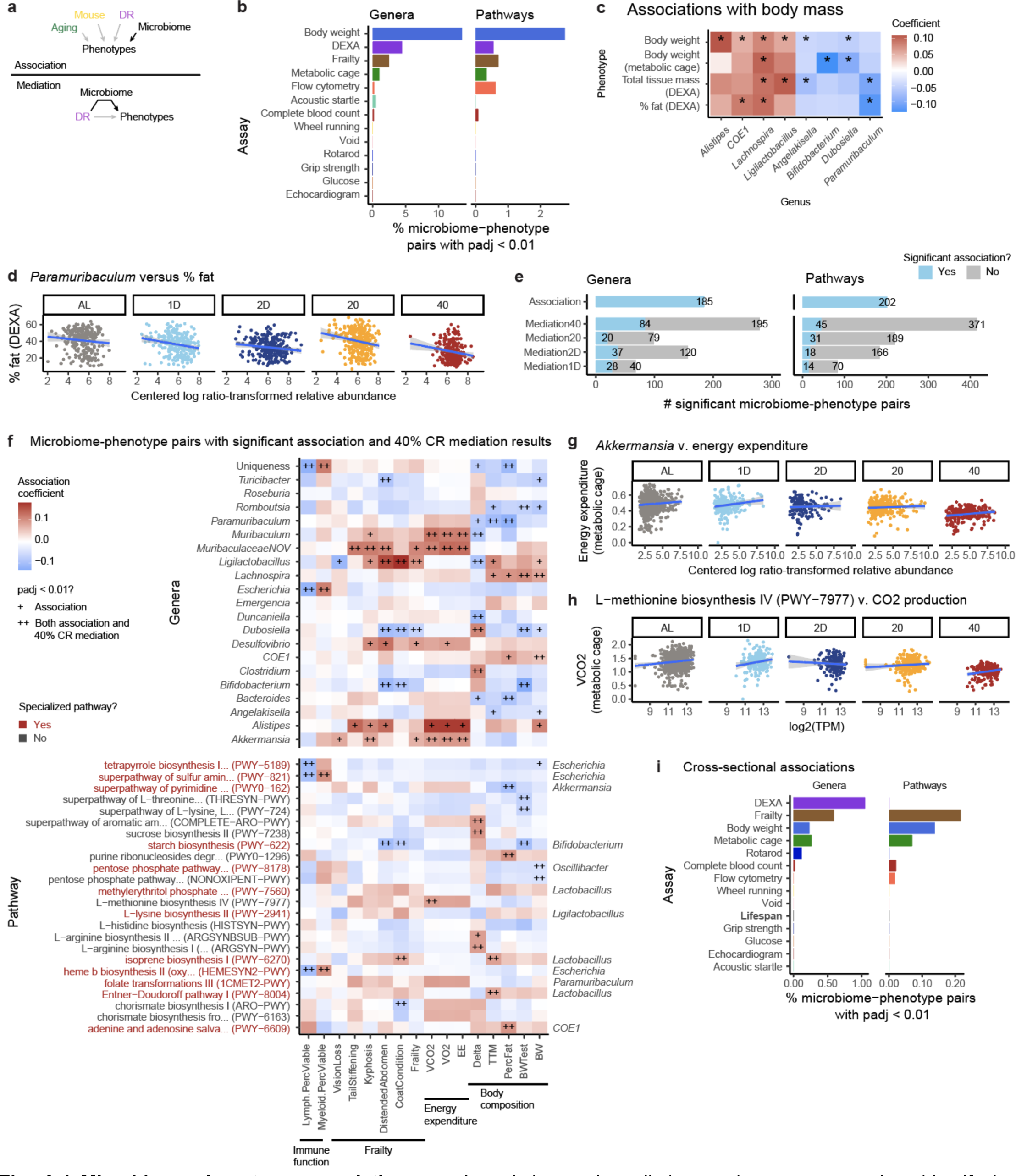
**Microbiome-phenotype associations. a**, Association and mediation analyses were used to identify host phenotypes influenced by the microbiome. For association analysis, a linear mixed model was fit for every microbiome feature-phenotype pair, with age, DR, and mouse as covariates. For mediation analysis, we tested each microbiome feature-phenotype pair to see which effects of DR are mediated by the microbiome. **b**, Percentage of significant (adjusted p-value < 0.01) microbiome-phenotype associations per phenotypic assay. The denominator for the percentage is the number of microbiome-phenotype pairs tested within each assay. **c**, Select associations between genera and body mass-related phenotypes. Positive associations are red, negative associations are blue. * indicates adjusted p-value < 0.01. **d**, *Paramuriculum* is associated with percent fat, as measured by dual-energy X-ray absorptiometry (DEXA). Blue line shows line of best fit and 95% confidence interval. **e**, Overlap of microbiome-phenotype pairs with significant (adjusted p-value < 0.01) association and mediation results. **f**, Heatmap of select microbiome-phenotype pairs significant by just association analysis (indicated with +) or by both association and 40% CR mediation analysis (indicated with ++). Phenotypes are grouped by phenotypic domain (e.g., energy expenditure or frailty). Microbiome features are sorted alphabetically. Color indicates the association coefficient. Positive associations are red, negative associations are blue. Pathways labeled in red are specialized pathways, and their most similar genus is labeled on the right side of the heatmap. **g**, *Akkermansia* is associated with energy expenditure, as measured by metabolic cages. **h**, Methionine biosynthesis (PWY-7977) is positively associated with the volume of carbon dioxide produced, as measured by metabolic cages. **i**, Percentage of significant (adjusted p-value < 0.01) cross-sectional microbiome-phenotype associations per phenotypic assay. The denominator for the percentage includes associations across all five ages tested. Lifespan is bolded to emphasize the absence of significant microbiome-lifespan associations.

Focusing on phenotypes related to body mass and composition, we identified that genera positively associated with body mass included *Alistipes*, *COE1* (Lachnospiraceae family), *Lachnospira,* and *Ligilactobacillus*, while genera negatively associated with body mass included *Angelakisella*, *Bifidobacterium*, *Dubosiella*, *and Paramuribaculum* (**Fig. 6c, d**).

To determine whether the microbiome was involved in the impact of DR on host phenotypes, we used mediation analysis (**Fig. 6a**, see Methods). Several hundred microbiome-phenotype pairs were significant (mediation effect adjusted p-value < 0.01) by mediation analysis, and a subset of these overlapped with significant associations (**Fig. 6e**). The mediation effect was small (mean proportion mediated = 19% for both genera and pathways), consistent with the microbiome having a modulatory, rather than driving, role in mediating the effect of dietary restriction on host physiology.

We focused on the microbiome-phenotype pairs with a significant association and significant 40% CR mediation result (**Fig. 6f**). These microbiome-phenotype pairs represent hypotheses about causal microbiome effects on host physiology. For example, *Muribaculum*, a novel genus in the Muribaculaceae family, and *Akkermansia* (**Fig. 6g**) are positively associated with energy expenditure. Indeed, the association between *Akkermansia* and increased energy expenditure has been reported previously^54^. Furthermore, pathway-phenotype associations propose mechanisms for the taxonomic associations. For example, the methionine biosynthesis pathway (PWY- 7977) is associated with increased carbon dioxide production — a parameter used to calculate energy expenditure — suggesting that the methionine may be involved in the associations with energy expenditure.

Lastly, we asked whether the microbiome is associated with lifespan. Because lifespan is not a longitudinal measurement, we performed association analysis separately per age (see Methods). Consistent with the longitudinal analysis (**Fig. 6b**), we identified numerous associations with frailty and body composition (**Fig. 6i**) but no significant associations with lifespan, indicating that several parameters of host health are affected by the microbiome, but that lifespan is not a microbiome-associated trait.

## Discussion

We generated nearly 3000 metagenomic profiles from more than 900 genetically diverse mice to conduct a comprehensive analysis of factors shaping the microbiome and how the microbiome itself influences host physiology. We integrated our dataset with host genomes, with a second longitudinal mouse metagenomic dataset, with thousands of human metagenomic samples, and with hundreds of longitudinal host phenotypes. Our analysis generated four major insights.

First, while microbiome information can be used to predict host age, we find that age-associated microbiome changes are better explained by cumulative exposure to stochastic environmental influences (neutral theory) than by the influence of the host (selection theory). This finding is consistent with a recent publication arguing that increased stochasticity is the signal used by methylation clocks to successfully predict an animal’s age^55^. We propose that microbiome uniqueness is a suitable proxy for quantifying the microbiome’s cumulative exposure to stochastic events.

Second, we find that host genetics explain similar amounts of microbiome variation as aging and dietary restriction, and that most microbiome features show significant but modest heritability. We demonstrate that detecting this genetic influence requires not only large sample sizes but also a controlled laboratory environment, such that the effect of host genetics is not obscured by unexplained environmental variability. Notably, even in a controlled study like DRiDO, the overall effect of unexplained environmental variability is larger than the effect of all other measured variables combined (**Fig. 4c**).

Third, we find that dietary restriction induces an older-appearing microbiome. We speculate that this finding is consistent with the age of the microbiome reflecting its cumulative exposure to stochastic events. The microbiomes of mice on dietary interventions were exposed to a diversity of feeding conditions — ranging from a 48-hour fasting state to a state of having just rapidly consumed three days’ worth of food — while the microbiomes of AL mice only experienced the fed state. We posit that this diversity of intestinal states increases the microbiome’s cumulative exposure to stochastic events that in turn affect its composition and function.

Lastly, we find that the microbiome is associated with aspects of host health — namely, body composition, immune function, and frailty — but not with longevity. In other words, we do not see evidence for the microbiome mediating the lifespan-extending effect of dietary restriction.

In summary, this work advances our understanding of the factors shaping the microbiome and elucidates which aspects of host physiology are influenced by the microbiome.

## Supplementary Discussion

### Aging

An important question addressed by our study is whether the gut microbiome contains a signature of host age. We discovered a large number of age-associated microbiome changes, but the only taxonomic change that was consistent across two mouse cohorts and thousands of human samples was uniqueness.

Wilmanski and colleagues first described uniqueness increasing with age in humans^26^, but this trend had not yet been observed in model organisms. By recapitulating this phenomenon in mice living under strictly controlled housing conditions, we provide evidence that this pattern is not due to confounding variables inherent to human studies — such as different diets, living environments, or medications — but that it may be a general feature of aging gut microbiomes.

Wilmanski and colleagues further argued that uniqueness characterizes not only aging, but specifically healthy aging. Like Ghosh and colleagues^25^, we did not find support for this second argument: uniqueness was not associated with frailty or lifespan, and its negative association with the proportion of lymphocytes (**Fig. 6**) is reminiscent of myeloid skewing, a detrimental phenomenon associated with aging^56^.

Rather, we propose that uniqueness quantifies the microbiome’s cumulative exposure to stochastic events. In addition to increasing with age, we show that uniqueness is increased by dietary restriction and, through the cohousing experiment, that uniqueness remains high in young mice harboring an old microbiome (**Fig. 3j**). Aging increases the microbiome’s exposure to stochastic events through the passage of time, DR increases exposure by subjecting the microbiome to a variety of fed and fasted intestinal states, and the cohousing experiment shows that an old microbiome in a young mouse retains its history of stochastic events.

More broadly, the cohousing experiment suggests that age-associated microbiome changes are better explained by cumulative exposure to stochastic events (neutral theory) than by the influence of the host (selection theory). Arguments have been made for^49^ and against^48^ the utility of neutral theory in describing the gut microbiome. We do not claim that selection theory has no applicability to the gut microbiome, but rather that age-associated changes, particularly in young mice, are better described by neutral theory.

Besides uniqueness, we identified several age-associated microbial pathways, including decreased production of vitamins like thiamine and riboflavin. An age-associated decrease in microbial production of thiamine may help to explain why older individuals have an increased prevalence of thiamine deficiency^57–59^. Moreover, age-associated decreases in vitamin biosynthesis pathways have been reported previously in mice^15^. It remains to be seen whether these functional changes are also the result of accumulating stochastic events or caused by an aging host.

### Genetics

In line with previous studies^27,28^, we find that the proportion of variance explained by host genetics is small (mean 7% across genera and 5% across pathways), and the majority of variance remains unexplained (**Fig. 4c**). However, we also find that many microbiome features (66% of genera and 51% of pathways) are significantly heritable, and the influence of genetics is as large as that of aging and dietary restriction.

We argue that we observed higher heritability than typically reported in human studies for two reasons: 1) large sample size and 2) limited environmental variability due to a controlled laboratory environment. Downsampling our dataset from thousands of samples to hundreds of samples erased heritability (**Extended Data Fig. 6**d). The necessity of large sample sizes for detecting microbiome heritability has also been reported previously^34,60^. Grieneisen and colleagues further argued that longitudinal data was critical for detecting microbiome heritability^34^, but we find that large sample size, rather than longitudinal data *per se*, is more important (**Extended Data Fig. 6**d). We note that Bruijning and colleagues have pointed out that large sample sizes alone may lead to spuriously high estimates of microbiome heritability when using relative abundances^60^. We attempted to avoid this issue by using centered log ratio-transformed data.

We believe the second reason why we observed higher heritability than expected is related to environmental variability. Heritability is defined as VG / (VG + VE), where VG is the amount of variance explained by additive genetic effects, and VE is all other sources of variance. We posit that VE — unexplained environmental variability — is higher in observational human studies than in studies involving agricultural animals and higher still than in studies with laboratory animals, and that larger VE leads to lower heritability estimates (**Fig. 4b**). An exception to this model is the extremely high microbiome heritability reported by Grieneisen and colleagues in baboons^34^. We believe the explanation for this exception is that Grieneisen and colleagues included a plethora of covariates in their linear modeling, like seasonality and dietary composition, that allowed them to effectively decrease the amount of unexplained environmental variance VE.

Despite widespread heritability, we were unable to find any QTLs that were consistent across ages. One explanation for this paradox is that we identified age-specific host-microbiome interactions. We recently demonstrated in this same cohort of DO mice that the influence of genetics on body weight varies over time and in different environments^61^, so it is plausible that the microbiome is also influenced by temporally variable genetic effects. Another explanation is that the genetic influence on the microbiome is extremely polygenic, and there are no individual loci that drive the association (i.e., our age-specific QTLs were false positives). Distinguishing between these explanations will be left for future investigations. QTL mapping has been performed previously in DO mice but never at multiple ages^50,62^.

### Dietary restriction

We found that both caloric restriction and fasting strongly modulate the microbiome. Perhaps unsurprisingly, the intensity of DR is proportional to the magnitude of microbiome changes (**Fig. 5f**), with 40% CR causing the largest changes. Overall, the effects of CR and fasting were concordant (**Fig. 5g**), but they induced slightly more distinct taxonomic changes than functional changes, suggesting that different types of DR lead to more varied compositional changes than functional changes.

We were surprised to find that DR induced an older-appearing microbiome state, in contrast to prior reports^51,63,64^. This conclusion was supported by two different analyses: a random forest classifier predicted older host age for DR samples than for AL samples, and PCoA plots showed DR and aging shifting the microbiome in the same direction. As discussed above, we propose that the apparent age of the microbiome as well as uniqueness reflect the microbiome’s cumulative exposure to stochastic events, not the biological age of the host or the chronological age of the microbiome. Therefore, an “older-appearing” microbiome state should not be interpreted as DR increasing host biological age, but rather as DR subjecting the microbiome to a greater diversity of intestinal conditions. The literature provides examples for both beneficial^11^ and detrimental^65^ consequences of microbiome adaptations to DR.

### Associations with host phenotypes

Consistent with the well-known link between obesity and the microbiome^35–37^, we observed numerous microbiome associations with body mass and composition. In addition, we observed associations with immune parameters, hematological parameters, and measures of frailty. Mediation analysis demonstrated that individual microbiome features are involved in some of the effects of DR on host physiology, but the proportion of the total effect mediated by the microbiome is small, and the assumption of unidirectional mediation is likely not fulfilled by microbiome data. Taken together, these results indicate that the microbiome modulates, rather than drives, the effect of DR on several aspects of host physiology.

In contrast, we saw no associations with lifespan. A lack of association with lifespan is not surprising; given that the microbiome undergoes even daily fluctuations, it seems improbable that a snapshot of the microbiome could predict mortality years later. Furthermore, the lack of a microbiome-lifespan association is consistent with our observation that DR induces an “older-appearing” microbiome state; even though DR clearly extends lifespan (**Fig. 1b**), this effect is independent of the gut microbiome. In the companion manuscript by Di Francesco et al., we demonstrate that other host phenotypes are predictive of lifespan, just not microbiome features.

### Resource

Several aspects of the DRiDO microbiome study make it a unique resource to the research community. First, it is the largest-to-date mouse microbiome dataset. The data are longitudinal and, because we performed metagenomic sequencing, provide both taxonomic and functional information. Second, we performed thorough and conservative quality control, including a pipeline to confirm that stool samples unambiguously matched their corresponding host genomes (**Extended Data Fig. 2**, **Supplementary Note 1**). Third, microbiome measurements are paired with host genomes and hundreds of longitudinally-collected host phenotypes. In addition to the raw sequencing files, we have made available summarized data tables, code, and an example analysis tutorial (https://github.com/levlitichev/DRiDO_microbiome) to facilitate use of this dataset by the community.

## Supplementary Note 1: Identifying sample mix-ups

### 1. Introduction

Sample mix-ups and well-to-well contamination are common in microbiome studies^66^. In this study, we were able to identify and remedy potential mix-ups or contamination due to our unique experimental design. Because every Diversity Outbred mouse was genotyped, we could compare each mouse genome to the small fraction of host reads in each stool sample to confirm that every stool sample came from the expected mouse (**Extended Data Fig. 2**a). To do this, we implemented a previously published pipeline^38^ as a Sunbeam^67^ extension: https://github.com/levlitichev/sbx_mbmixture. Below, we refer to our implementation of this pipeline as “mbmix”.

### 2. Genotyping

Mice were genotyped with the GigaMUGA genotyping array, which contains 143,259 markers. The genomes of DO mice contain large regions inherited directly from one of eight founder lines, so we are able to impute many more genomic positions than those directly measured by the genotyping array. After quality control, 946 genotypes remained. For stool samples belonging to mice without a genotype, we were unable to apply mbmix.

### 3. Description of method

For every sample, we filtered to mouse reads by mapping quality-controlled reads with bwa^68^ to the mouse genome (mm10). An average of 1.1 M reads (9.1% of quality-controlled reads) could be mapped to the mouse genome. These mouse reads were compared to every mouse genotype. At each single nucleotide polymorphism (SNP), we counted the number of times that a read agreed or disagreed with the expected nucleotide. Summing across all SNPs, we reported the discordance between a stool sample and a mouse genotype as the proportion of discordant SNPs.

An example of concordance is shown in **Extended Data Fig. 2**b: the stool sample from mouse DO-AL-0020 at 97 weeks of age has the lowest proportion of mismatches with the genotype of mouse DO-AL-0020. Therefore, we are confident that this stool sample came from the expected mouse. An example of discordance is shown in **Extended Data Fig. 2**c: the stool sample from mouse DO-20-1188 at 21 weeks of age has much lower discordance with mouse DO-2D-4188, rather than the expected mouse DO-20-1188. In this case, the likely explanation is that somewhere during data generation — probably during stool collection or DNA extraction — “20” was mistaken for “2D,” and the wrong tube was picked up. This is an example of a stool sample that was discarded because we could not confidently determine from which mouse it originated. For more details about this method, please see Lobo et al. 2021.

### 4. Overview of sample mix-ups

mbmix was run on all samples. The primary outcome of the pipeline was the proportion of discordant SNPs against each mouse genome. Samples with considerably fewer discordant SNPs against a different genome than against the expected genome (self_proportion_mismatch - best_proportion_mismatch > 0.05) were considered potential sample mix-ups. During our manual inspection, we also assessed whether mix-ups were geographically concentrated in certain regions of 96-well plates.

After extensive manual inspection of 4352 sequenced samples, we discarded or renamed 886 samples (20.3%, **Extended Data Fig. 2**d**, Supplementary Table 1**). Samples were renamed when we could confidently determine which stool sample was actually sequenced. We now address each of the reasons why samples were discarded or renamed.

#### 4.1 Suspicious region, cause unknown

The largest category of potential sample mix-ups consisted of samples in suspicious geographic regions of 96- well plates (**Extended Data Fig. 2**e). Specific examples of suspicious geographic regions are listed in **Supplementary Table 2**. In certain cases, the cause of the mix-ups could be traced to a specific step of data generation (for example, tubes that were dropped during DNA extraction of samples on final plate 27), but for most cases, it was unclear whether the mix-ups occurred during stool collection, DNA extraction, or library preparation.

#### 4.2 Within-cage

The next largest category of potential mix-ups were geographically scattered samples whose predicted mouse was in the same cage. Within-cage mix-ups were particularly likely to occur because samples from the same cage were handled together (during stool collection, DNA extraction, and library preparation), but another equally likely explanation is coprophagy, i.e., mice consuming each other’s feces. We are unable to distinguish between these two possibilities, so we took the conservative approach of discarding these samples.

#### 4.3 Index collision

We identified one clear example of an index collision, where one library batch (LB46) accidentally received the same sequencing indexes as another library batch (LB83). This mistake required discarding a full 96-well plate of samples plus a smaller number of samples that were sequenced as part of a redo sequencing run. Because the DNA from these samples was unaffected, we were able to successfully remake and sequence libraries from the DNA.

#### 4.4 Suspicious region, cause known

These samples correspond to a geographic region of potential mix-ups where we were able to confidently determine what stool sample was actually sequenced (**Supplementary Table 2**). For example, on final plate 7, one column of samples all corresponded to the mouse in the adjacent column. This likely represented a pipetting error during library preparation.

#### 4.5 Cross-cage

These samples correspond to scattered samples in which the predicted mouse was not in the same cage as the expected mouse. These mix-ups are very unlikely to be explained by coprophagy. Possible explanations include well-to-well contamination or sample mishandling during DNA extraction or library preparation. These samples were discarded.

#### 4.6 Very low uniqueness

These samples had a borderline mbmix result (self_proportion_mismatch - best_proportion_mismatch < 0.05, but > 0) and high similarity to another microbiome sample. These samples were identified by calculating all pairwise distances between microbiome profiles and subsetting to samples frequently involved in very small pairwise distances (< 0.15). For each of these “very low uniqueness” samples, we asked whether the most similar microbiome profile belonged to the same mouse as predicted by mbmix. We identified 15 samples where this was the case. For example, the most similar sample to DO_2D_4051_069w was DO_2D_4050_069w, and the predicted mouse by mbmix was DO-2D-4050, suggesting that DO_2D_4051_069w was either a duplicate of or contaminated by DO_2D_4050_069w. These 15 “low uniqueness” samples were discarded.

#### 4.7 Animal swap

Lastly, mbmix identified one instance where two mice were accidentally put back into the wrong cages after a phenotyping session and misidentified the rest of their lives. Microbiome profiles from mice AL-0097 and AL- 0105 correctly matched their corresponding genotypes at 44 weeks of age, but afterwards, samples from mouse

AL-0097 appeared to come from AL-0105, and samples from AL-0105 appeared to come from AL-0097 (**Extended Data Fig. 2**f). Both mice were white, had the same ear tag, and were housed in adjacent cages. Examining the longitudinal body weight for these two mice revealed that they were swapped at approximately 56 weeks (**Extended Data Fig. 2**g). Microbiome samples from these mice after 44 weeks were renamed, and the body weight data was fixed.

### 5. Conclusion & discussion

The mbmix pipeline allowed us to identify and remedy errors made during data generation in order to increase the quality of our final dataset. We discarded 775 (17.8%) and renamed 111 (2.5%) samples. In the previous application of this pipeline, 22 of 297 (7.4%) samples were discarded^38^. Why did we discard a larger fraction of samples? First, we applied this pipeline on cohoused mice. As a result, there were 255 samples where we were unable to distinguish between a potential sample mix-up and coprophagy. We made the conservative decision to discard these samples. Second, we discarded 136 samples because of one index collision. Excluding these two issues, our proportion of discarded samples (9.7%) more closely resembles that of Lobo et al. We suspect that this range of sample mix-ups (5-10%) is common in large-scale microbiome sequencing studies, but few experimental designs allow for identifying and fixing these errors.

## Methods

### DRiDO study

The Dietary Restriction in Diversity Outbred mice (DRiDO) study is described in depth in Di Francesco et al. Briefly, 960 DO mice were enrolled in quarterly waves from breeding generations 22-24 and 26-28. Just one female mouse was used from each litter, so no mice in the study were siblings. Mice were housed up to eight animals per pen. Only female mice were used to prevent aggressive competition for limited food resources^69^.

At six months of age, mice began one of five dietary interventions: *ad libitum,* 20% caloric restriction, 40% caloric restriction, one day per week fasting, or two consecutive days per week fasting. 960 mice were enrolled in the study, but only 937 mice were alive at six months. Dietary restriction was initiated at six months to evaluate the consequences of adult-onset, rather than lifelong, DR. 20% CR mice received 2.75 grams of food per day, 40% CR mice received 2.06 grams per day, 1D mice were fasted from Wednesday 3pm until Thursday 3pm, and 2D mice were fasted from Wednesday 3pm until Friday 3pm. CR mice received a triple feeding on Friday afternoon. They consumed this food quickly, causing the 20% CR and 40% CR mice to undergo approximately one day or two days of fasting, respectively, over the weekend.

Body weight was measured weekly, while a variety of other phenotypes were collected every six months or yearly (**Extended Data Table 1**). The experiment was conducted at The Jackson Laboratory (Bar Harbor, ME), and animal procedures were approved by the Animal Care and Use Committee at The Jackson Laboratory (protocol # 06005).

### Stool collection

Mice were scruffed, and one stool pellet was collected fresh. The pellet was added to RNAlater and stored at - 80°C.

### DNA extraction

Stool pellets were removed from RNAlater with clean tweezers. DNA was extracted with the QIAGEN DNeasy PowerSoil Pro Kit (Cat. # 47016) following manufacturer instructions with one modification: pellets were homogenized for one minute using the MP Biomedicals FastPrep-24 Classic Bead Beating Grinder and Lysis System, rather than using a vortex adapter. DNA was eluted in 50 uL of C6. DNA concentration was measured using a NanoDrop Spectrophotometer. DNA samples with concentration < 10 ng/uL were excluded from further data generation. DNA was extracted across 124 batches of 24 or 48 samples. We included at least one negative control (nothing added to lysis tube at start of protocol) and one positive control every ∼100 samples. We used two different positive controls: ATCC 10 Strain Even Mix Whole Cell Material (Cat. # MSA-2003) or ZymoBIOMICS Gut Microbiome Standard (Cat. # D6331). These controls contain cells from 10 and 21 microbes, respectively, with a variety of relative abundances.

### Library preparation

Extracted DNA was prepared for sequencing using the Illumina DNA Prep Kit (Cat. # 20060059). We added 7.5 uL of input DNA diluted to 3-22 ng/uL. We modified the protocol to use ¼ of the manufacturer recommended reagent volumes. Samples were barcoded using IDT for Illumina DNA/RNA UD Indexes Sets A and B (Cat. # 20027213, 20027214), except for a handful of samples that were barcoded with Nextera DNA CD Indexes (Cat. # 20018708). Samples were eluted in 35 uL of elution buffer. DNA concentration was measured using the Qubit dsDNA High Sensitivity Assay Kit (Cat. # Q32851). If library preparation was unsuccessful (i.e., concentration < 1 ng/uL), we attempted to redo the library starting from the input DNA. We examined a random subset of samples from each batch on a TapeStation 4200 instrument using a High Sensitivity D1000 ScreenTape (Cat. # 5067-5584). Library preparation was performed in 96-well plates across 70 batches of 48 or 96 samples each. We included at least one negative control (7.5 uL of dilution buffer added to well at start of protocol) and one positive control every ∼100 samples. The positive control was the ATCC 10 Strain Staggered Mix Genomic Material (Cat. # MSA-1001), which contains DNA from 10 bacteria at a variety of relative abundances.

### Pooling and sequencing

Libraries were pooled and sequenced on a NovaSeq 6000 with paired ends (2x150 bp). Sequencing was performed across eight sequencing runs and two NovaSeq 6000 machines: one at Calico Life Sciences LLC and one at the University of Pennsylvania.

### Data preprocessing

We performed quality control of our sequencing reads using the Snakemake pipeline Sunbeam^67^ (v2.1.1). More specifically, we removed adapters with cutadapt^70^ (v3.1, forward and reverse adapters = CTGTCTCTTATACACATCT); we trimmed low-quality bases and discarded low-quality reads with trimmomatic^71^ (v0.39, ILLUMINACLIP:trimmomatic/adapters/NexteraPE-PE.fa:2:30:10:8:true LEADING:3 TRAILING:3 SLIDINGWINDOW:4:15 MINLEN:36); we discarded reads with many repetitive sequences with komplexity (https://github.com/eclarke/komplexity); and we removed host reads (mean 9% of input reads) using bwa^68^ (v0.7.17) against the mm10 genome.

### Quality control

We collected 4214 stool samples from 944 DO mice (16 mice died prior to the first collection). Of these 4214 stool samples, 3586 stool samples (85%) were sequenced. We were unable to sequence 628 stool samples because either A) we could not successfully extract DNA (the DNA concentration had to be > 10 ng/uL) or B) we could not prepare a library from the DNA (the library concentration had to be > 1 ng/uL) despite multiple attempts. After accounting for positive and negative controls and some stool samples being sequenced more than once, we sequenced 4352 samples.

Samples were discarded for any of the following reasons (in this order):

1. The sample did not definitively come from the expected mouse (see “Identifying sample mix-ups” below), n=775
2. We discovered that the date of sample collection was after the animal’s date of death, so we could not be certain that the stool sample was correctly identified, n=6
3. The sample received too few reads (< 750k read pairs for stool samples, < 100k read pairs for positive controls), n=29
4. The proportion of reads assigned to the mouse genome was suspiciously high (> 50%), n=53
5. The sample was very different from all other samples (see “Discarding outliers” below), n=13

After these quality-control steps, we were left with 3473 samples, corresponding to 2997 stool samples (71% of the original 4214) from 913 DO mice.

### Identifying sample mix-ups

Because every DO mouse was genotyped, we could compare the small fraction (∼9%) of host reads in each stool sample to each mouse genome to confirm that every stool sample came from the expected mouse. We did this by adapting a previously published pipeline^5^. The idea is to count the number of times that host reads are discordant with the host genotype. After running this pipeline on all samples, we discarded 775 samples and renamed 111 samples. Notably, 136 of the discarded samples were due to one mistake (an index collision between two library batches), and 225 of the discarded samples may not have been mix-ups but the result of coprophagy. The small number of renamed samples involved situations in which we could confidently identify the cause of the mistake. For complete details, please see **Supplementary Note 1**.

### Discarding outliers

We calculated all pairwise sample distances to identify potential outliers (**Extended Data Fig. 3**a). The largest distances were enriched for 13 samples: the 45,348 pairwise distances greater than 0.9 involved 3310 samples, but just 13 samples were involved in 48.3% of these distances. These 13 samples were considered outliers and omitted from analysis.

### Taxonomic and functional classification

After quality control, we performed taxonomic and functional classification with two distinct approaches. First, we used HUMAnN3 (ref. ^41^) for taxonomic (MetaPhlAn4, ref ^72^) and functional profiling with the following databases: mpa_vOct22_CHOCOPhlAnSGB_202212, full_chocophlan.v201901_v31, and uniref90_annotated_v201901b. We used all default parameters and added ‘--unclassified-estimation’ to estimate the proportion of unclassified reads. We also performed taxonomic profiling with a second approach: Kraken2 (ref. ^39^, v2.1.2) with the Mouse Gastrointestinal Bacterial Catalogue^40^ (MGBC) and default parameters. Because the Kraken2+MGBC approach classified more reads (**Extended Data Fig. 3**b) and returned fewer uncharacterized taxa, we present Kraken taxonomic results except in two places: positive control stacked barplots (because MGBC does not contain all microbes present in the positive controls, **Extended Data Fig. 1**c) and comparisons to the other datasets (see “Comparison to B6 and human datasets” below).

We identified 376 genera and 482 species using MetaPhlAn, 252 genera and 1093 species with Kraken, and 422 MetaCyc^42^ pathways with HUMAnN3. Because so many microbes in the mouse gut microbiome are uncharacterized, we focused on genera rather than species for better interpretability.

To account for differences in sequencing depth, we calculated genus-level relative abundances (exclusive of unclassified reads), and we divided pathway reads-per-kilobase (RPK) abundances by the sample sum (excluding UNMAPPED and UNINTEGRATED) and multiplied by 1 million. This is equivalent to the transcripts-per-million (TPM) normalization used in RNA sequencing^73^, so we adopt this nomenclature even though we are not referring to transcripts.

Samples produced from the same stool sample were aggregated together. For Kraken taxonomic results, we aggregated by summing absolute counts. For HUMAnN results, we aggregated by taking the mean genus-level relative abundance and mean pathway TPM abundance.

We distinguished between “community-wide” and “specialized” pathways on the basis of their similarity to taxonomic features. Specialized pathways were defined as having a small number (1-4) of Pearson correlations with genera above 0.5. Correlations were calculated using centered log ratio-transformed relative abundances for genera and transcripts-per-million (TPM) abundances for pathways.

### Data normalization prior to linear modeling

For downstream linear modeling, we used centered log ratio-transformed, genus-level relative abundances and pathway TPM abundances after replacing zeros. Zeros were replaced with each feature’s minimum non-zero value divided by two.

We excluded low prevalence features. For Kraken taxonomic results, we considered the 100 most abundant genera (based on total counts across all samples), which accounted for >99% of all genus-level counts. For HUMAnN, we retained pathways with at least 100 TPM in at least 10% of samples (273 of 422 pathways).

We included several community features in our linear modeling: three genus-level measures of ɑ-diversity (Shannon index, Simpson index, and Chao1 index), the first three principal coordinates (based on Bray-Curtis distance of genus-level relative abundances), taxonomic uniqueness (also based on genus-level Bray-Curtis distance), and functional uniqueness (based on Euclidean distance of log2(TPM) pathway abundances). Uniqueness is defined as the distance (or β-diversity) of a microbiome sample to its nearest neighbor^26^.

Finally, prior to use in a linear model, features were scaled so that estimated coefficients were comparable across features.

### PCoA and PCA

Principal coordinate analysis (PCoA) plots were based on Bray-Curtis distances of genus-level relative abundances. The principal component plot (PCA) plot was based on Euclidean distance of pathway log2(TPM) abundances.

### PERMANOVA

We performed permutational multivariate analysis of variance (PERMANOVA) using the adonis2 function from the vegan^74^ R package with the following parameters: formula = dist ∼ age + DR, by = “margin”, permutations = 999. Age is a continuous variable encoding age in months, DR is a categorical variable encoding each of the 5 dietary interventions (samples collected prior to dietary randomization are considered AL). We ran PERMANOVA on all 2997 samples. For genera, we used Bray-Curtis distance on relative abundances. For pathways, we used Euclidean distance on log2(TPM) abundances.

### Linear mixed model per microbiome feature

To assess the influence of age, dietary restriction (DR), and genetics on each microbiome feature (y_mb), we fit the following linear mixed model:

1. y_mb ∼ age + DR + (1|mouse) + (1|cohort) + (1|batch) + (1|cage) + (1|genetics)

Where age (in weeks, scaled) is a fixed effect; DR is a fixed effect with five levels (AL, 1D, 2D, 20, 40); (1|mouse) is a random intercept; (1|cohort) is a random intercept corresponding to DO breeding cohorts; (1|batch) is a random intercept corresponding to microbiome DNA extraction batch; (1|cage) is a random intercept corresponding to the cage in which a mouse was housed for the entirety of its life; and (1|genetics) is a random effect corresponding to additive genetic effects, which we encode by providing the kinship matrix. In ecology, this model is referred to as the repeated measures animal model^75^. The (1|mouse) random effect accounts for “repeatability” or “permanent environment effects”, while (1|genetics) accounts strictly for additive genetic effects.

We used the previously published^61^ kinship matrix for this cohort of DO mice. Importantly, we multiplied the kinship matrix (in which the diagonal was approximately 0.5) by 2 before using it in the linear mixed model. This step is necessary because the genetic covariance matrix in a linear mixed model must contain coefficients of relationships, which are twice the kinship coefficients^76^.

The significance of each fixed effect was evaluated using a conditional Wald test. The significance of each random effect was evaluated using a likelihood ratio test where the null model omitted the random effect. P-values were adjusted with the Benjamini-Hochberg procedure, separately for taxonomic and functional features. Adjusted p-values < 0.01 were considered significant.

Model 1 was fit using ASReml-R^77^ (v4.1.0.716), as well as lme4qtl^78^ (v0.2.2) to confirm identical results with a different software package (**Extended Data Fig. 6**b).

### Age prediction

We used a random forest classifier (randomForest R package v4.7-1.1 with default parameters) to predict the age of a mouse based on its microbiome profile. We performed age prediction in 3 different contexts: 1) predicting the age of DO AL mice, 2) predicting the age of DR mice using a classifier trained on AL mice, and vice versa; and 3) predicting age in the cohousing experiment.

For predicting age of DO AL mice, we split into training (70%) and testing (30%) sets while stratifying by cage, i.e., samples from the same cage were not present in both the training and testing set. We used out-of-bag predictions when reporting training accuracy. The classifier did not see samples from the testing set except when making its final prediction. Age was treated as a continuous variable. We used relative abundances for the top 100 most abundant genera and log2(TPM) abundances for the 272 pathways that passed prevalence filtration.

For predicting age of DR mice, the classifier was trained on all AL samples and evaluated on all DR samples. We used out-of-bag predictions when reporting AL accuracy. For predicting age of AL mice, we trained a classifier on all 40% CR samples and evaluated it on all other samples, including the other DR groups. Age was treated as a continuous variable, and we again considered 100 genera and 272 pathways.

For predicting age of mice in the cohousing experiment, the classifier was trained on all baseline samples and tested on all other samples. Age was treated as a binary variable (young or old). We considered relative abundances for all 125 genera (see “Cohousing experiment”).

### Longitudinal B6 mouse cohort

Fifteen four-week-old male C57BL/6 (“B6”) mice were ordered from The Jackson Laboratory. Mice were housed five per cage across three cages. Stool pellets were collected upon arrival and then every three months. Pellets were collected fresh into empty 1.7 mL tubes and frozen at -80°C. Animal procedures were approved by the Institutional Animal Care and Use Committee at the Perelman School of Medicine at the University of Pennsylvania (protocol # 806361).

DNA was extracted with the Qiagen Dneasy PowerSoil Pro Kit (Cat. # 47016), libraries were prepared with the Nextera DNA Flex Library Prep Kit (Cat. # 20018705), and libraries were sequenced on a NextSeq 550 machine with 75-bp single-end sequencing. Data were analyzed the same way as the DO samples: pre-processed with Sunbeam and then taxonomically and functionally classified with HUMAnN.

### Human metagenomic data

Human metagenomic sequencing data was obtained using the curatedMetagenomicData^46^ package (v3.6.2). We filtered for stool samples from individuals meeting the following criteria: age ≥ 18 years, “healthy” or “control”, and no current antibiotic use. Furthermore, we only included studies with ≥ 50 individuals meeting these criteria and an age interquartile range ≥ 5 (to make sure each study had a diversity of ages). This resulted in 4101 individuals from 20 studies. We obtained genus-level relative abundances and pathway RPK abundances, which we normalized to log2(relative abundances) and log2(TPM) values after replacing zeros, as described above.

### Comparison to B6 and human datasets

We considered only samples from AL mice when comparing the DO cohort to the B6 cohort and human samples. We fit the following linear models to identify age-associated microbiome features in each dataset:

2. DO AL mice: y_mb ∼ age + (1|mouse) + (1|cohort) + (1|batch) + (1|cage)

3. B6 mice: y_mb ∼ age + (1|cage)

4. Humans: y_mb ∼ age + (1|study)

Where (1|study) is a random intercept corresponding to one of 20 studies comprising the human cohort. See “Linear mixed model per microbiome feature” for details about the other terms. Models 2, 3, and 4 were fit with MaAsLin2 (ref. ^79^, v1.12.0).

Because the human dataset had been processed with HUMAnN, we used our MetaPhlAn taxonomic results for the DO AL mice, instead of the Kraken taxonomic results. We used log2-transformed genus-level relative abundances after zero replacement. For prevalence filtration, we retained genera with at least 0.001% relative abundance in at least 10% of samples (DO AL: 248 of 311, B6: 262 of 310, humans: 90 of 331) and pathways with at least 100 TPM in at least 10% of samples (DO AL: 262 of 399, B6: 233 of 329, humans: 358 of 573).

We also tested several measures of ɑ-diversity (Shannon index, Simpson index, inverse Simpson index, and richness) and uniqueness. For DO AL mice, uniqueness was recalculated considering just AL mice. For humans, uniqueness was calculated separately within each of 20 studies.

The significance of the age coefficient was evaluated using a conditional Wald test. P-values were adjusted with the Benjamini-Hochberg procedure, separately per dataset and separately for taxonomic and functional features. Due to the additional burden of needing to be consistent across datasets, the adjusted p-value threshold was increased to 0.1 for this analysis.

For select microbiome features, we also fit the following basic linear model separately within each human study to assess consistency across studies (**Extended Data Fig. 5**a-c):

5. Within each human study: y_mb ∼ age

The 20 p-values (one from each human study) were adjusted with the Benjamini-Hochberg procedure.

### Cohousing experiment

We obtained n=25 8-week-old female C57BL/6 mice from The Jackson Laboratory and n=20 19- and 20-month-old female C57BL/6 mice from the National Institutes on Aging. Prior to the start of the experiment, mice were housed five per cage with other mice of the same age. Mice were allowed to acclimate for at least one week prior to the start of the experiment. During cohousing, three young mice were housed with two old mice for one month. Control young and old mice remained housed with other mice of the same age. After one month, two of the cohousing cages were separated by age and two cohousing cages remained cohoused. Stool pellets were collected at baseline, after one month of cohousing, and after two, four, six, and eight weeks of separation. Animal procedures were approved by the Institutional Animal Care and Use Committee at the Perelman School of Medicine at the University of Pennsylvania (protocol # 806361).

We performed 16S sequencing of these stool samples. DNA was extracted using the Qiagen DNeasy PowerSoil Pro Kit (Cat. # 47016). We amplified the V1/V2 variable region using KAPA HiFi HotStart ReadyMix (Roche, Cat. # KK2602) and the 27F/338R primer pair (27F: 5’-TATGGTAATTGTAGAGTTTGATCCTGGCTCAG-3’, 338R:

5’-XXXXXXXXXXXXTGCTGCCTCCCGTAGGAGT-3’, where the sequence of X’s represents the sample-specific barcode). We performed a bead-based clean-up of the pooled libraries using AMPure XP SPRI beads (Cat. # A63881). Libraries were paired-end (2x250 bp) sequenced across two MiSeq runs. The first run contained samples collected at baseline, at 4 weeks of cohousing, and at 4 weeks of separation. The second run contained samples collected at 2, 6, and 8 weeks of separation. Data were processed using QIIME2 (ref. ^80^, v2023.2.0). Reads were demultiplexed^81^ and denoised with DADA2 (ref. ^82^). For the first run, the forward read was trimmed to 250 bp and the reverse read to 165 bp. For the second run, the forward read was trimmed to 240 bp and the reverse read to 220 bp. Reads from the two runs were then merged and taxonomically classified against the SILVA 138 database^83,84^ using a naïve Bayes classifier^85,86^.

### Heritability

Using model 1, heritability was calculated as the variance assigned to the (1|genetics) random effect divided by total variance. The standard error of heritability was estimated using the vpredict function from ASReml. A feature was considered heritable if it had an adjusted p-value < 0.01 (likelihood ratio test, followed by Benjamini-Hochberg for p-value adjustment).

We also calculated heritability with the following cross-sectional model, separately at each of 5, 10, 16, 22, and 28 months (**Extended Data Fig. 6**d):

6. y_mb ∼ DR + (1|cohort) + (1|batch) + (1|cage) + (1|genetics)

The DR term was omitted from the 5-month analysis because this was prior to dietary randomization. Model 6 was fit with ASReml. For this cross-sectional analysis, heritability was computed using all samples at a given age. We also fit model 1 after downsampling to 110 mice in order to obtain a similar number of samples as in cross-sectional data (**Extended Data Fig. 6**d).

### Comparison to heritability estimates from other studies

For **Extended Data Fig. 6**c, we compared our heritability estimates to those reported in Supplementary Table 3A by Schlamp and colleagues^50^. This table contained 27 operational taxonomic units (OTUs) that could be classified to the level of genus. Of these 27, only 8 overlapped with the genera that we tested for heritability. For **Fig. 4b**, we plotted the proportion of significantly heritable taxa (based on the authors’ definitions) from 8 other _studies_29–32,50,87–89.

### Comparing all experimental variables

To compare the effects of all experimental variables to each other, we used a modified version of model 1 in which age and DR were encoded as random intercepts:

7. y_mb ∼ (1|age) + (1|DR) + (1|mouse) + (1|cohort) + (1|batch) + (1|cage) + (1|genetics)

This allowed us to compare the variance explained by each variable. Age was provided as a categorical variable. p-values were adjusted separately for each experimental variable. Model 7 was fit with ASReml.

### Quantitative trait loci (QTL) mapping

We performed QTL mapping as described previously^90^. Briefly, we tested the association between each SNP marker against each of 107 microbiome features (top 100 genera plus seven community features) at each of 5 ages: 5, 10, 16, 22, and 28 months. Dietary group and DO mouse cohort were included as covariates, except at 5 months (prior to the start of DR) when only cohort was included as a covariate. QTL mapping was performed with R/qtl2 (ref. ^91^) using previously published leave-one-chromosome-out kinship matrices and genotype probabilities^90^. p-values were calculated based on permutation and adjusted using the Benjamini-Hochberg procedure.

### Prediction of dietary group

We used a random forest classifier to predict the dietary group of DO mice. We performed prediction separately per age. Within each age, we performed 10-fold cross-validation while stratifying by cage so that no samples from the same cage were present in both the training and validation set. We used predictions on the validation sets when reporting accuracy. Dietary group was treated as a categorical variable, and we considered 100 genera and 272 pathways.

### Microbiome-phenotype associations

To identity associations between microbiome features and host phenotypes, we tested all microbiome-phenotype pairs with the following model:

8. y_pheno ∼ y_mb + age + DR + (1|mouse)

Where y_pheno is a phenotypic trait such as body weight, age and DR are fixed effects, and (1|mouse) is a random intercept. For each phenotype, we selected the measurement closest in time to each microbiome sample and only included microbiome-phenotype pairs obtained within 100 days of each other. The significance of each microbiome-phenotype association was calculated using a likelihood ratio test where the null model omitted y_mb. We tested 100 genera and 252 pathways against 197 phenotypic traits measured across 13 assays.

Model 8 could not be used to test for association for lifespan (a non-longitudinal measurement), so we fit the following cross-sectional model at each of 5, 10, 16, 22, and 28 months:

9. y_pheno ∼ y_mb + DR

The DR term was omitted from the 5-month analysis because this was prior to dietary randomization. Model 8 was fit with lme4 (ref. ^92^, v1.1-33) while model 9 was fit with the lm function in R.

### Mediation

Mediation analysis estimates the proportion of treatment T’s effect on outcome Y that is mediated by mediator M. In our study, the treatment T is dietary restriction, the outcome Y is a phenotypic trait, and the mediator M is a microbiome feature. We performed mediation analysis using the model-based approach within the mediate R package^93^ (v4.5.0). We fit the following two models:

10. Mediator model: y_mb ∼ DR_X + age + (1|mouse)

11. Outcome model: y_pheno ∼ y_mb + DR_X + age + (1|mouse)

Where DR_X corresponds to one of the 4 dietary interventions. Mediation analysis was performed separately for each of the DR groups. We report the average causal mediation effect (ACME) estimate and p-value. P-values were adjusted with Benjamini-Hochberg separately per dietary group. Microbiome-phenotype pairs with an ACME adjusted p-value < 0.01 were considered significant. We tested the same features as for association analysis. When reporting the mean proportion mediated, we consider only significant mediation results where the direct effect and mediation effects estimates have the same sign.

### Statistics

p-values were adjusted using the Benjamini-Hochberg procedure. Significance was defined as an adjusted p-value < 0.01, unless explicitly stated otherwise. t-tests were used for pairwise comparisons. All t-tests were two-sided except in **Fig. 5h**. Tests were unpaired except in **Fig. 5f**. For boxplots, boxes extend from the 25^th^ to 75^th^ percentiles, whiskers extend to 1.5 times the interquartile range, and the center line is the median. p-value symbols are defined as follows: ns : p ≥ 0.05, * : p < 0.05, ** : p < 0.01, *** : p < 0.001, **** : p < 0.0001. Downstream analysis and plotting was performed in RStudio (R v4.2) using the tidyverse^94^ (v2.0.0). phyloseq^95^ (v1.42.0), and vegan^74^ (v2.6.4) packages. Final figures were created with Adobe Illustrator.

### Code and data availability

Raw sequencing fastq files and summarized data tables will be made available on SRA. Code for reproducing manuscript figures is available at https://github.com/levlitichev/DRiDO_microbiome. Host phenotypes collected as part of the DRiDO study are available at https://doi.org/10.6084/m9.figshare.24600255.v1. The genetic kinship matrix and genotype probabilities are available at https://doi.org/10.6084/m9.figshare.13190735.

## Acknowledgements

For their assistance with sequencing, we thank Margaret Roy and Twaritha Vijay at the Calico Sequencing Core, Jonathan Schug at the Penn Next-Generation Sequencing Core, and Ana M. Misic at Illumina, Inc. For helpful discussions and suggestions, we thank the members of the Thaiss and Levy labs.

## Author Contributions

Lev Litichevskiy: investigation, formal analysis, data curation, writing - original draft, writing - review & editing. Maya Considine: investigation. Jasleen Gill: investigation. Vasuprada Shandar: investigation. Timothy O. Cox: investigation. Hélène C. Descamps: investigation. Kevin M. Wright: software, formal analysis. Kevin R. Amses: software, formal analysis. Lenka Dohnalová: validation. Megan J. Liou: validation. Monika Tetlak: investigation. Mario R. Galindo-Fiallos: investigation. Andrea C. Wong: validation. Patrick Lundgren: validation. Junwon Kim: validation. Giulia T. Uhr: validation. Ryan J. Rahman: investigation. Sydney Mason: investigation. Carter Merenstein: formal analysis. Frederic D. Bushman: supervision. Anil Raj: software. Fiona Harding: formal analysis. Zhenghao Chen: formal analysis. G.V. Prateek: formal analysis. Martin Mullis: formal analysis. Andrew G. Deighan: data curation. Laura Robinson: investigation, data curation. Ceylan Tanes: software. Kyle Bittinger: supervision. Meenakshi Chakraborty: software. Ami S. Bhatt: supervision. Hongzhe Li: methodology, supervision. Ian Barnett: methodology. Emily R. Davenport: methodology. Karl W. Broman: methodology. Robert L. Cohen: conceptualization, funding acquisition. David Botstein: conceptualization, funding acquisition. Adam Freund: conceptualization, supervision, project administration. Andrea Di Francesco: supervision, project administration. Gary A. Churchill: formal analysis, data curation, supervision. Mingyao Li: methodology, supervision. Christoph A. Thaiss: supervision, funding acquisition, writing - original draft, writing - review & editing.

## Companion manuscript

Di Francesco, A., Deighan, A. G., Litichevskiy, L., Chen, Z., Luciano, A., Robinson, L., Garland, G., Donato, H., Schott, W., Wright, K. M., Raj, A., Prateek, G. V., Mullis, M., Hill, W., Zeidel, M., Peters, L., Harding, F., Botstein, D., Korstanje, R., Thaiss, C.A., Freund, A.F., Churchill, G. A. Regulators of health and lifespan extension in genetically diverse mice on dietary restriction.

## Extended Data Figures

**Extended Data Fig. 1.**
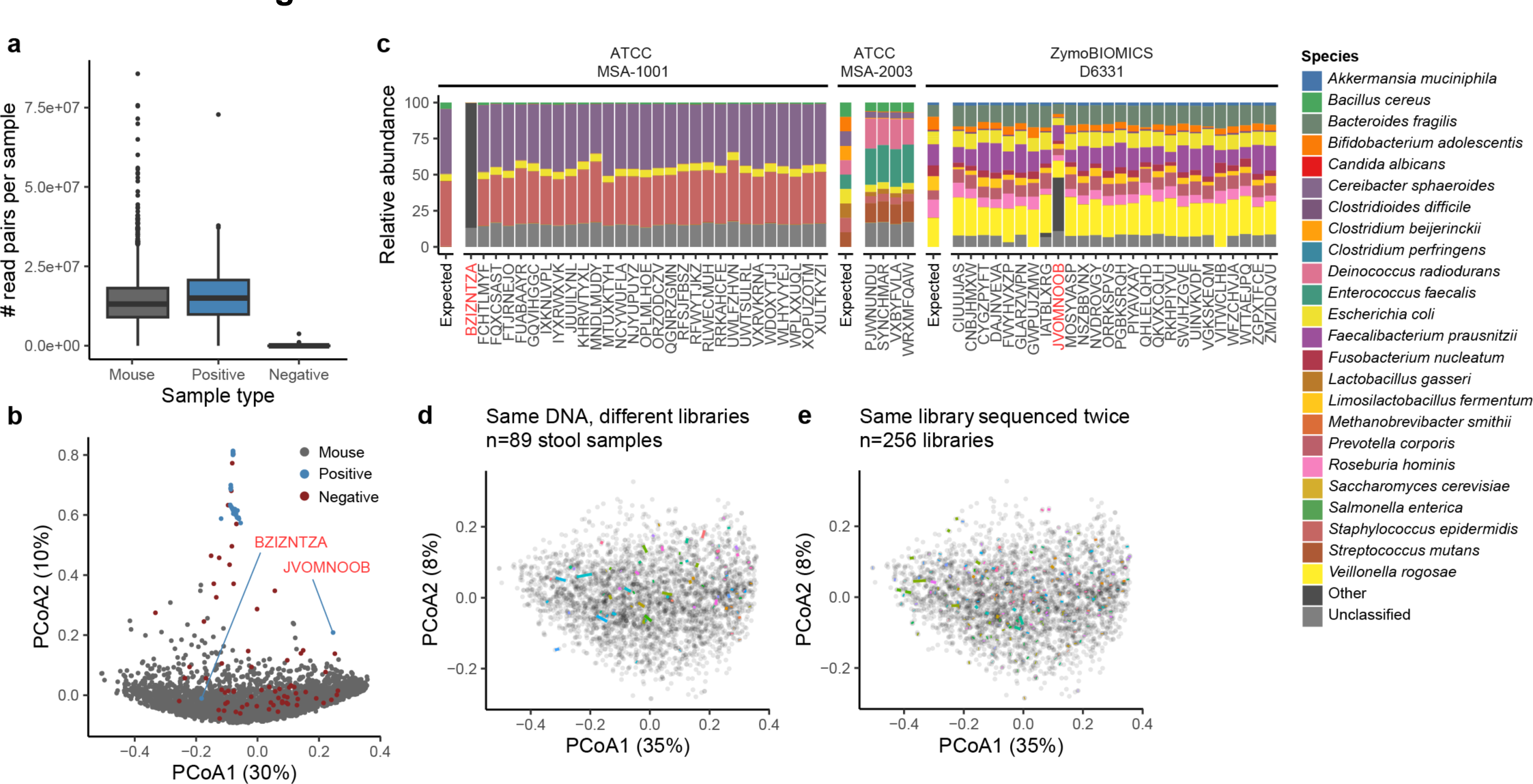
**Positive and negative controls. a**, Number of read pairs per sample (prior to aggregation), grouped by sample type. **b**, PCoA of all samples prior to aggregation. Two positive controls (BZIZNTZA and JVOMNOOB, highlighted in red) clustered separately from the other positive controls. PCoA1 and PCoA2 explain 30% and 10% of overall variance, respectively. **c**, Species-level relative abundances (MetaPhlAn4) for positive controls. Two positive controls (BZIZNTZA and JVOMNOOB, highlighted in red) did not display the expected community composition. **d**, PCoA of non-control samples prior to aggregation. Samples originating from the same DNA are connected by a colored line. **e,** Same PCoA plot as **d**, highlighting instances in which a library was sequenced multiple times.

**Extended Data Fig. 2.**
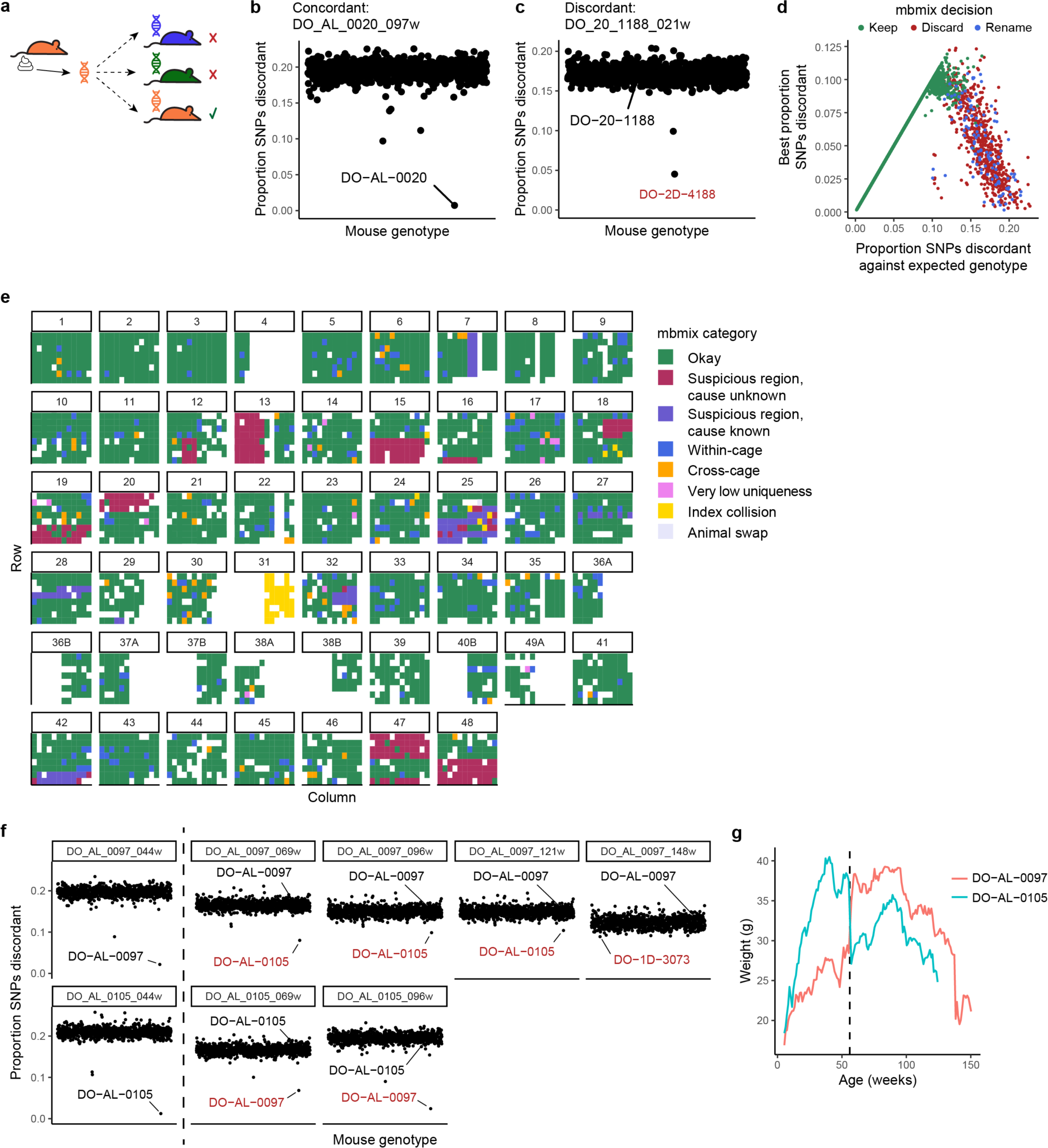
Identifying sample mix-ups. a, Sample mix-ups were identified by comparing host reads from each microbiome sample against all host genotypes (we term this pipeline “mbmix”). For more details, see **Supplementary Note 1**. **b,** Example of concordance between a microbiome sample and the expected host genotype. The x-axis is each host genotype, the y-axis is the proportion of single nucleotide polymorphisms (SNPs) that were discordant between the microbiome sample and the host genome. **c,** Example of discordance. Microbiome sample DO_20_1188_021w was supposed to originate from mouse DO-20-1188, but it appears to have come from DO-2D-4188. **d**, Best proportion discordant SNPs versus proportion of discordant SNPs against the expected genotype. The fate of each sample is indicated by its color: kept (green), discarded (red), or renamed (blue). **e**, Plate view of mbmix categorization. Each panel is a “final plate”, a 96-well plate of libraries prior to pooling. White regions either didn’t contain a sample, contained a sample that obtained no reads (e.g., left half of final plate 31), contained a sample whose mouse did not have a genotype, or contained a control sample. **f**, Proportion discordant SNPs for stool samples from mice DO- AL-0097 and DO-AL-0105. Samples from 44 weeks were concordant with the expected mouse genotype. All other samples from mouse DO-AL-0097 appeared to come from mouse DO-AL-0105, except DO_AL_0097_148w, which was inconclusive. The two other samples from mouse DO-AL-0105 appeared to come from DO-AL-0097. For discordant results, the mouse with the lowest proportion of discordant SNPs is colored red. **g**, Body weight for mice DO-AL-0097 and DO-AL-0105. The vertical dashed line at 56 weeks represents the likely time that these mice were swapped in their cages.

**Extended Data Fig. 3.**
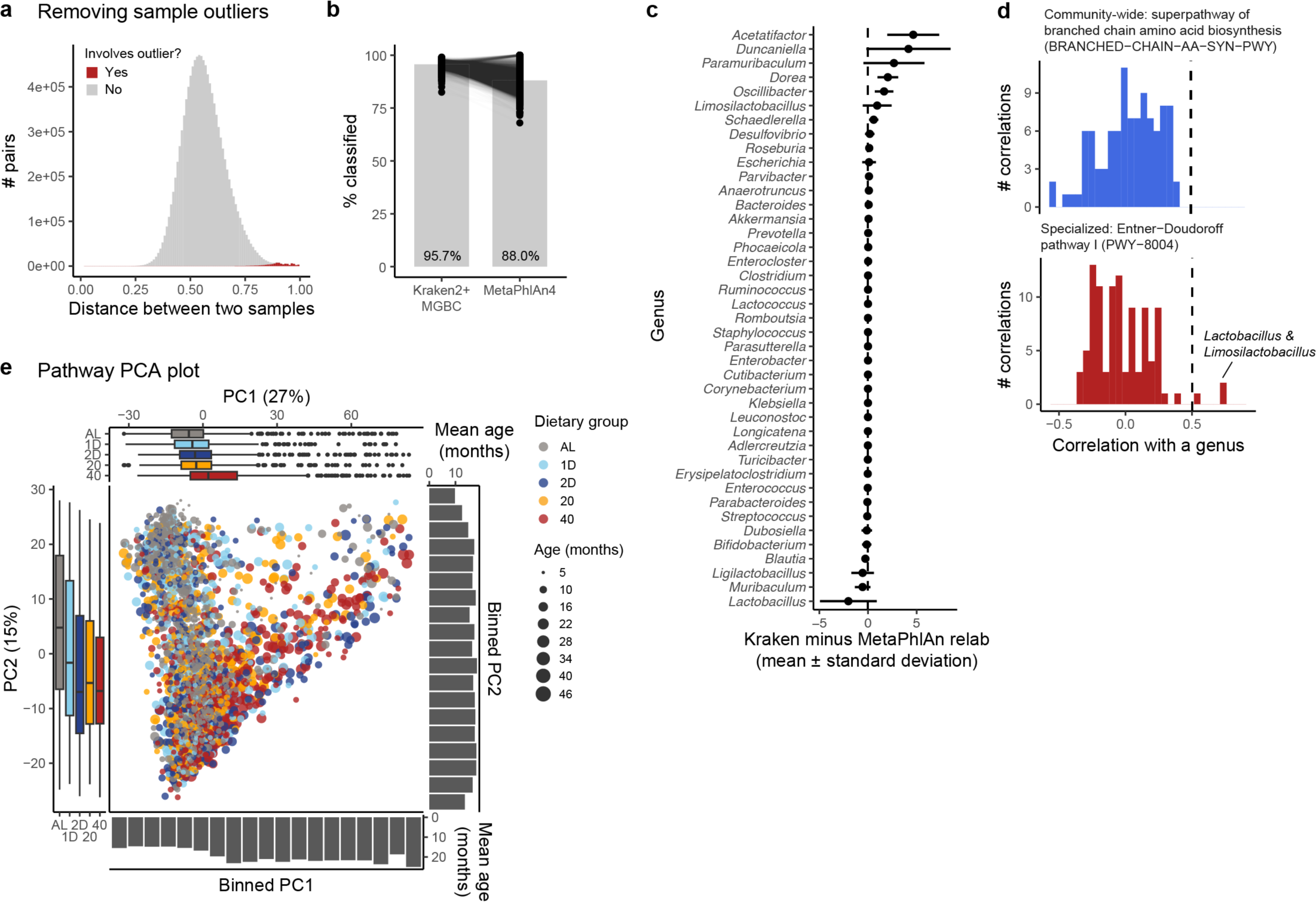
Additional quality-control and details related to taxonomic and functional classification. a, Histogram of all pairwise sample distances (Bray-Curtis on relative abundances). Distances involving any of 13 outlier samples are shown in red. **b**, Percentage of reads that could be classified for non-control samples using either Kraken2+MGBC or MetaPhlAn4. Mean percent classified indicated in black text. **c**, Difference between Kraken2+MGBC and MetaPhlAn4 genus-level relative abundances for the 41 genera present in both taxonomic databases. Each horizontal line shows the mean ± standard deviation across all non-control samples. **d**, Examples of community-wide and specialized pathways. The largest correlations for the specialized pathway (PWY-8004) were with *Lactobacillus* and *Limosilactobacillus*. **e,** PCA plot based on microbial pathways. PC1 and PC2 explain 21% and 8% of overall variance, respectively. For more details, see Fig. 1 legend.

**Extended Data Fig. 4.**
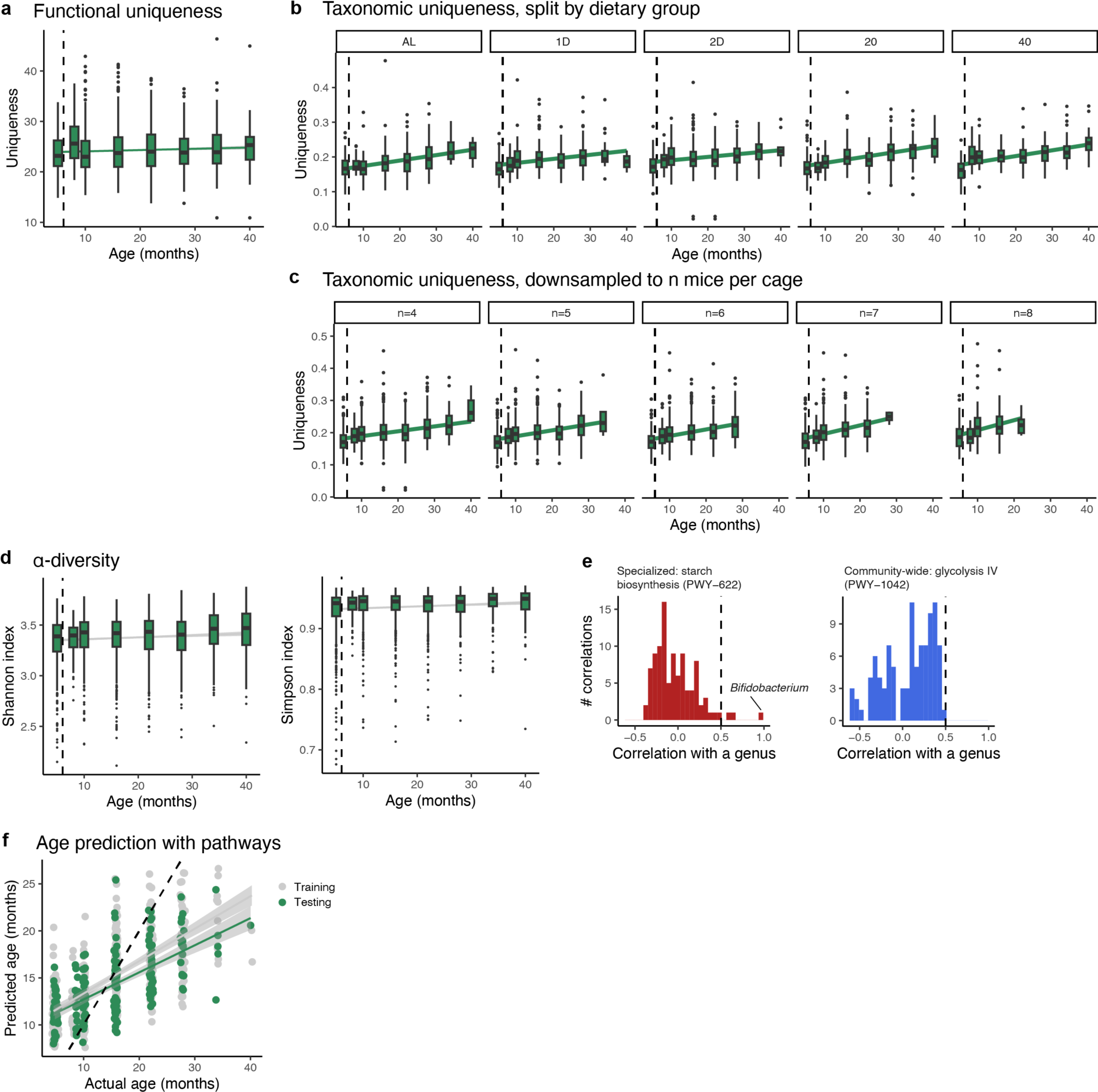
**Additional details related to age-associated microbiome changes in DO mice. a**, Functional uniqueness increases with age. Green line represents line of best fit and 95% confidence interval. Vertical dashed line at six months represents start of dietary restriction. **b**, Taxonomic uniqueness increases with age in all dietary groups. **c**, Taxonomic uniqueness increases with age even when the number of mice per cage is kept fixed. For various n, cages with at least n mice at that age were considered. If the number of mice was greater than n, then n mice were randomly chosen. Uniqueness was then recomputed on this subset of samples. **d**, ɑ-diversity (as measured by Shannon and Simpson indexes) has a non-significant (adjusted p-value > 0.01) positive association with host age. Line of best fit and 95% confidence interval shown in gray. **e**, Histograms of pathway-genus correlations. The largest genus correlation for the specialized pathway (PWY-622) is to *Bifidobacterium*. There are no correlations above 0.5 for the community-wide pathway (PWY- 1042). **f**, Age prediction based on pathways. The classifier was provided with pathway log2(TPM) profiles from AL mice. Gray and green dots are predictions on training and testing data, respectively. Black dotted line at y=x represents perfect accuracy.

**Extended Data Fig. 5.**
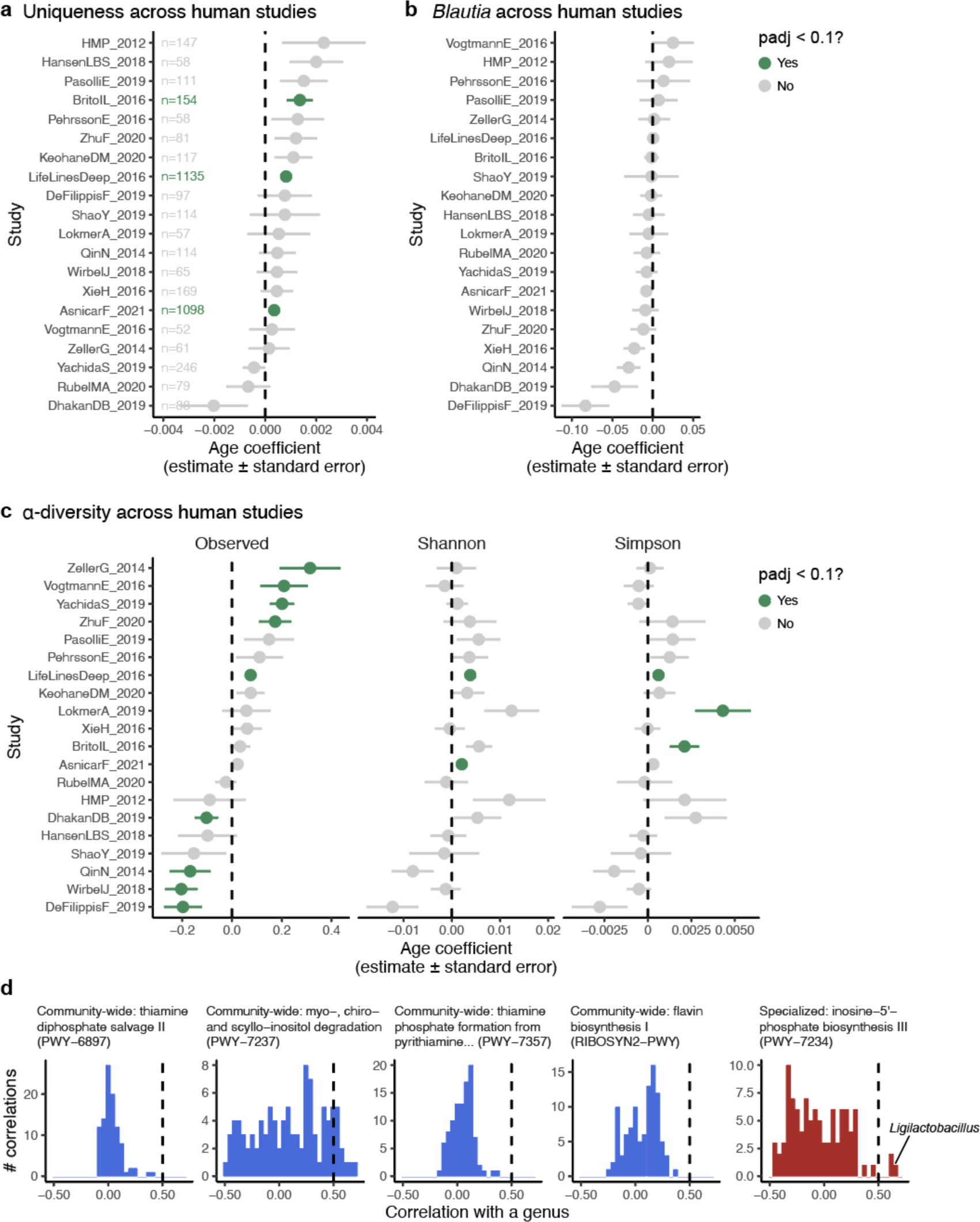
**Additional details related to universal age-associated microbiome changes. a**, Uniqueness increases with age in most human studies and significantly increases with age in the two largest studies. Adjusted p-values < 0.1 are shown in green. The number of individuals per study is indicated. **b**, *Blautia* decreases with age in just a few studies, and when regressing against age separately per study, no studies have an adjusted p-value < 0.1. **c**, ɑ-diversity versus age, separately per human study. p-values were adjusted separately per metric. **d**, Histograms of pathway-genus correlations. For the specialized pathway (PWY-7234), the largest genus correlation is to *Ligilactobacillus*.

**Extended Data Fig. 6.**
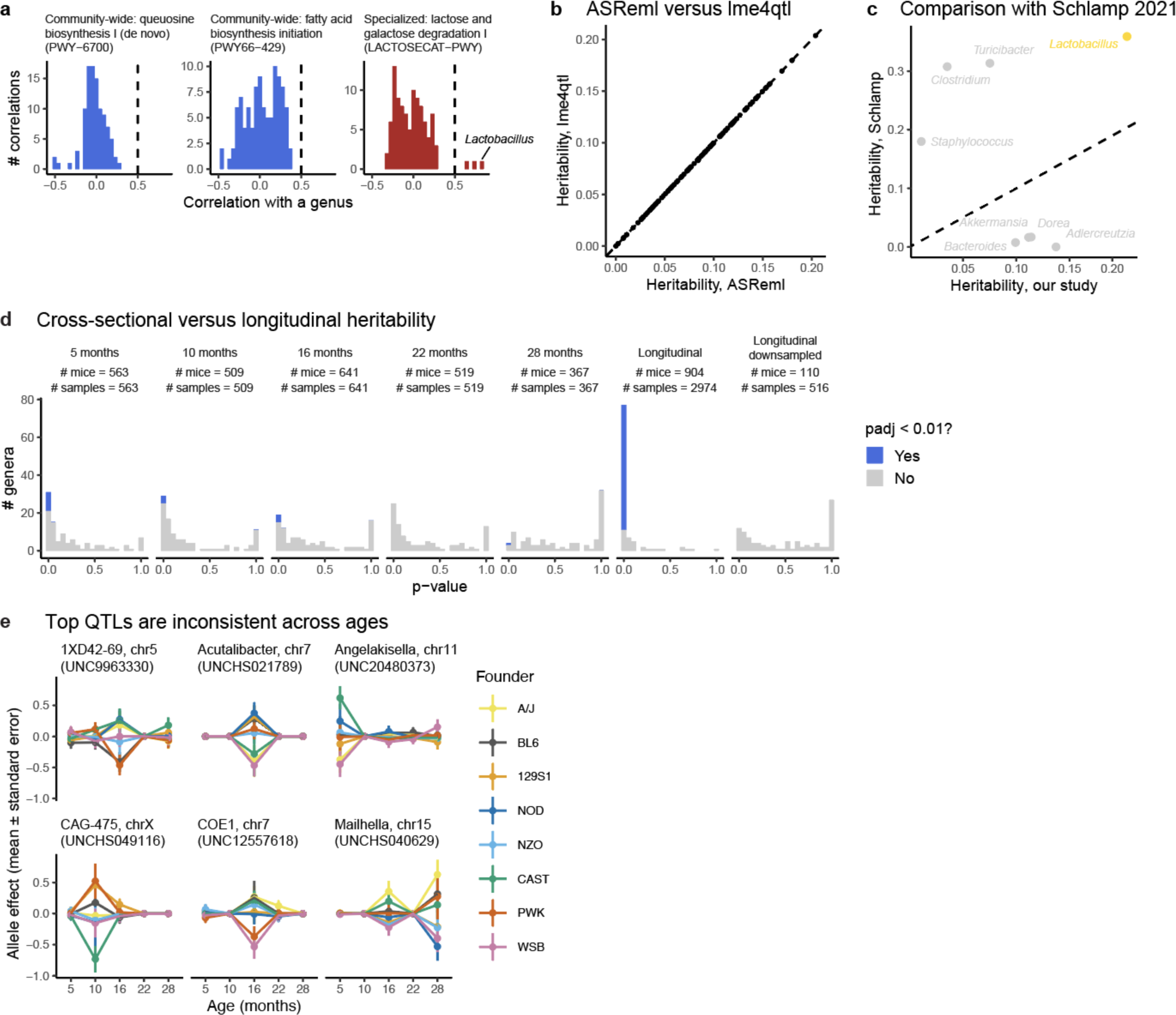
Additional details related to microbiome heritability. a, Histograms of pathway-genus correlations. For the specialized pathway (LACTOSECAT-PWY), the largest genus correlation is to *Lactobacillus*. **b**, Heritability computed with lme4qtl or ASReml using the same model and data. **c**, Comparison of heritability estimates from a different DO mouse study (Schlamp et al. 2021). Plot shows the eight genera for which heritability was assessed in both datasets. Of these eight, the most heritable taxon in both studies was *Lactobacillus* (highlighted in yellow). **d**, Cross-sectional versus longitudinal versus downsampled longitudinal heritability. Genera with significant heritability (adjusted p-value < 0.01) are shown in blue. The longitudinal results are the primary heritability results presented throughout the manuscript. **e**, Allele effects across ages for the top six age-specific QTLs (permutation test, adjusted p-value < 0.01). The title above each sub- panel indicates the genus, chromosome, and genotyping marker for the QTL result. Color of each line represents the allele effect for each of eight founders comprising the Diversity Outbred genetic pool.

**Extended Data Fig. 7.**
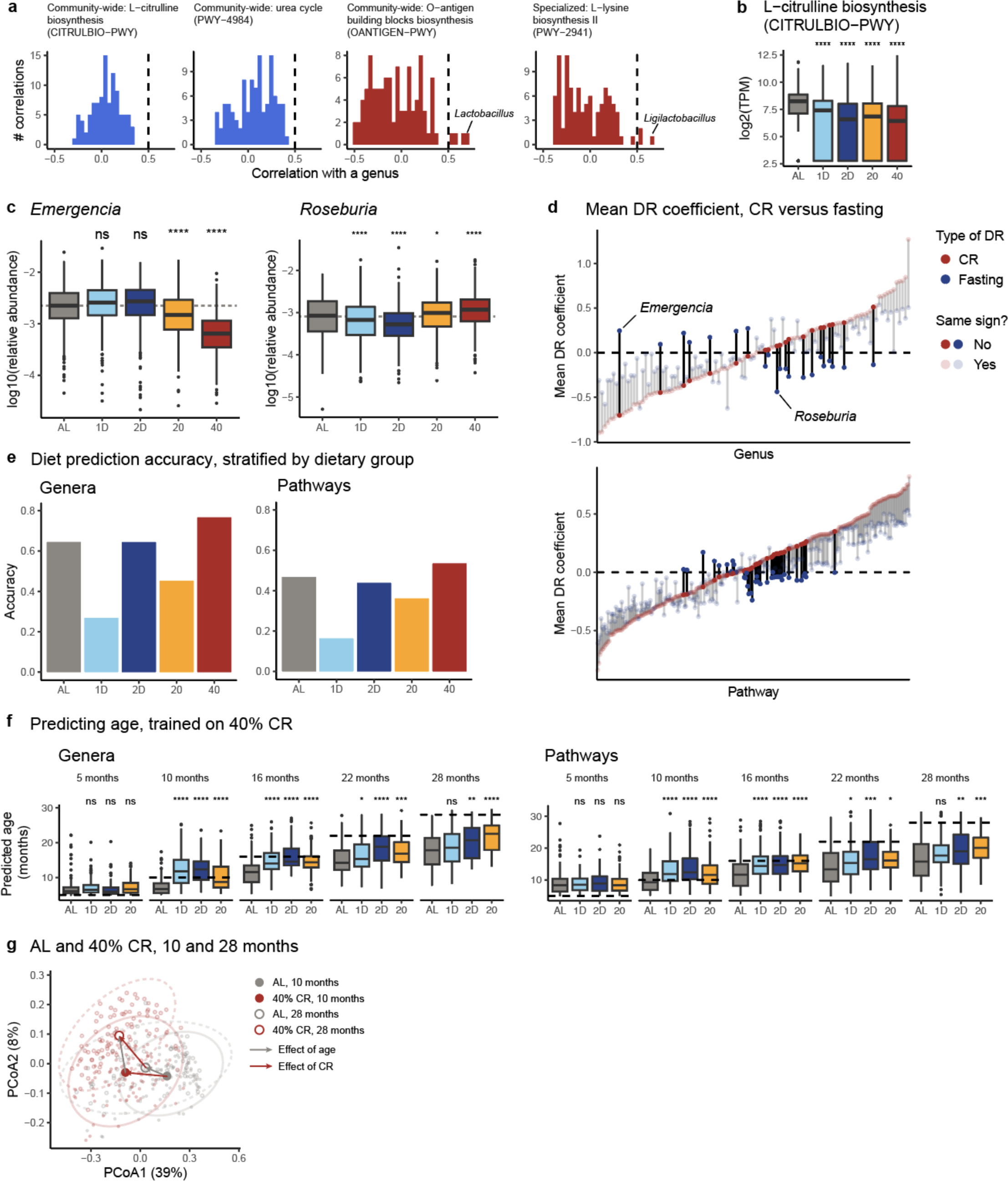
**Additional details related to the effects of dietary restriction on the microbiome. a**, Examples of community-wide and specialized pathways. The specialized pathways are most highly correlated with *Lactobacillus* and *Ligilactobacillus*. **b,** The L-citrulline biosynthesis pathway (CITRULBIO-PWY) is decreased by all DR groups. **c**, Taxonomic changes unique to fasting or caloric restriction. *Emergencia* is exclusively decreased by CR, while *Roseburia* is decreased by fasting and increased by caloric restriction. Horizontal dashed gray line at the AL group median is a visual aid to help compare across groups. Statistical significance evaluated with a t-test against the AL group. **d**, Mean CR versus mean fasting coefficients for genera (top) and pathways (bottom). Vertical lines highlight the difference in mean CR coefficients (red) versus mean fasting coefficients (blue). Features with opposite signs are opaque, while features with the same sign are transparent. Dashed horizontal line at 0. **e**, Diet prediction accuracy, stratified by dietary group, using genera (left) and pathways (right). Only predictions after the start of dietary restriction were considered. **f**, Age prediction of a classifier trained on all 40% CR samples and evaluated on all other samples, using genera (left) or pathways (right). Horizontal dotted line shows the actual age of samples collected at that timepoint. Statistical significance evaluated with a t-test. **g**, PCoA of AL and 40% CR samples from middle-aged (10 months) and old (28 months) samples. Ordination based on just these samples. Group centroids are depicted by the four large points, along with 95% data ellipses. Arrows connect group centroids to depict the effect of age (gray) and the effect of caloric restriction (red). PCoA1 and PCoA2 explain 39% and 8% of overall variance, respectively.

**Extended Data Fig. 8.**
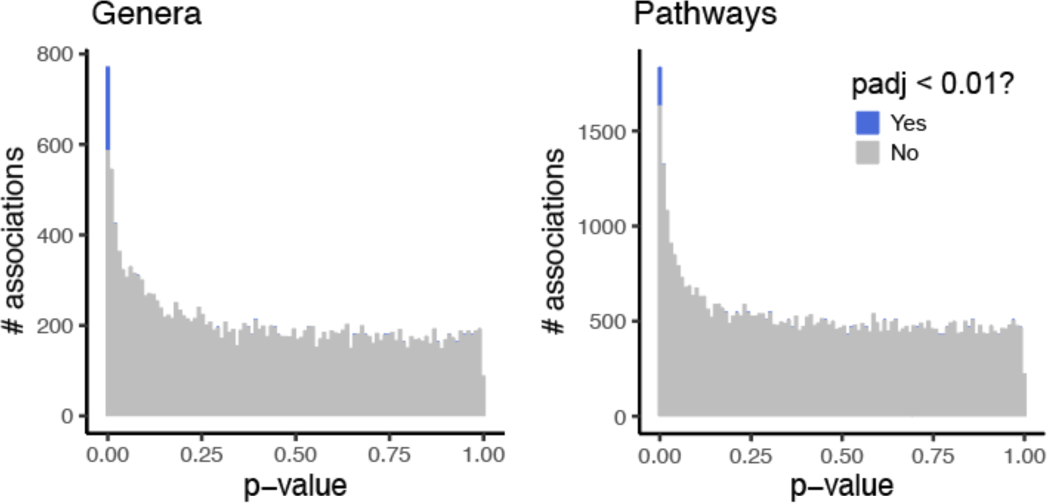
Additional details related to microbiome-phenotype associations. Histogram of p-values for associations between phenotypes and genera (left) or pathways (right). Associations with an adjusted p-value < 0.01 are shown in blue.

## Extended Data Tables

**Extended Data Table 1.**
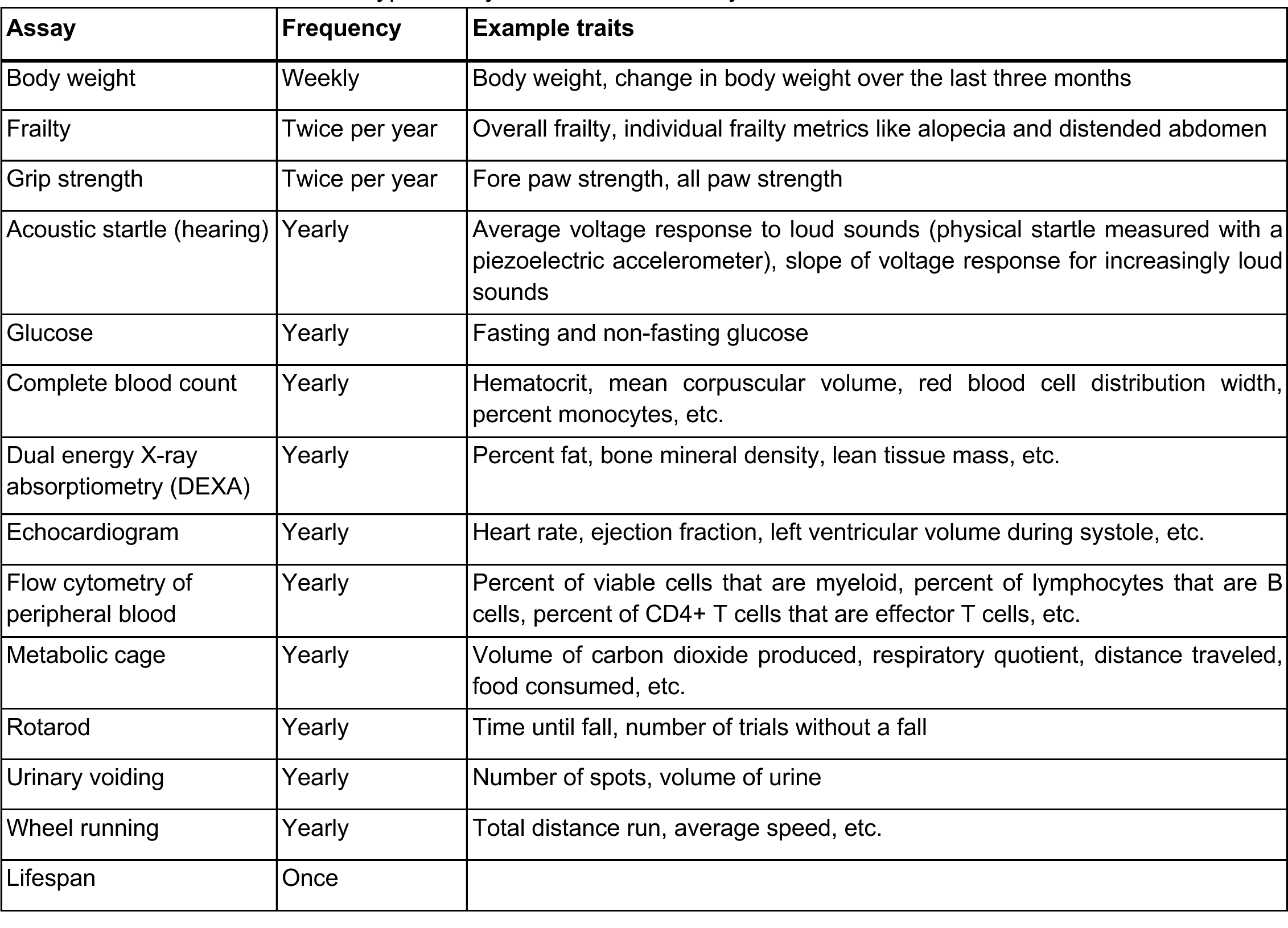
Phenotypic assays in the DRiDO study.

| Assay | Frequency | Example traits |
| --- | --- | --- |
| Body weight | Weekly | Body weight, change in body weight over the last three months |
| Frailty | Twice per year | Overall frailty, individual frailty metrics like alopecia and distended abdomen |
| Grip strength | Twice per year | Fore paw strength, all paw strength |
| Acoustic startle (hearing) | Yearly | Average voltage response to loud sounds (physical startle measured with a piezoelectric accelerometer), slope of voltage response for increasingly loud sounds |
| Glucose | Yearly | Fasting and non-fasting glucose |
| Complete blood count | Yearly | Hematocrit, mean corpuscular volume, red blood cell distribution width, percent monocytes, etc. |
| Dual energy X-ray absorptiometry (DEXA) | Yearly | Percent fat, bone mineral density, lean tissue mass, etc. |
| Echocardiogram | Yearly | Heart rate, ejection fraction, left ventricular volume during systole, etc. |
| Flow cytometry of peripheral blood | Yearly | Percent of viable cells that are myeloid, percent of lymphocytes that are B cells, percent of CD4+ T cells that are effector T cells, etc. |
| Metabolic cage | Yearly | Volume of carbon dioxide produced, respiratory quotient, distance traveled, food consumed, etc. |
| Rotarod | Yearly | Time until fall, number of trials without a fall |
| Urinary voiding | Yearly | Number of spots, volume of urine |
| Wheel running | Yearly | Total distance run, average speed, etc. |
| Lifespan | Once |  |

**Extended Data Table 2.** Significant (permutation test, adjusted p-value < 0.1) age-specific microbiome quantitative trait loci (QTLs).

| Microbiome feature | Age (months) | Chromosome | Position (bp) | LOD | p-value | Adjusted p-value |
| --- | --- | --- | --- | --- | --- | --- |
| Angelakisella | 5 | 11 | 116956666 | 8.33 | 0.01 | 0.078 |
| CAG-475 | 10 | X | 103262003 | 10.10 | 0.01 | 0.078 |
| 1XD42-69 | 16 | 5 | 115024909 | 8.40 | 0.01 | 0.078 |
| Acutalibacter | 16 | 7 | 135473480 | 8.42 | 0.01 | 0.078 |
| COE1 | 16 | 7 | 29431249 | 10.14 | 0.01 | 0.078 |
| Mailhella | 28 | 15 | 69199803 | 9.17 | 0.01 | 0.078 |

## Supplementary Tables

**Supplementary Table 1.** Summary of potential sample mix-ups identified by mbmix.

| Category | # samples | Action | Description |
| --- | --- | --- | --- |
| Suspicious region, cause unknown | 305 | discard | A region of a plate with many potential sample mix-ups, but the cause of the mix-ups is unknown. |
| Within-cage | 255 | discard | Scattered samples whose predicted mouse is in the same cage. Unknown if these are caused by coprophagy or errors during data generation. |
| Index collision | 136 | discard | LB46 received the wrong indexes, which caused it to collide with LB83. Prior to identifying the cause of the problem, these samples were re-sequenced as part of a redo run, and they again collided with other redo samples. |
| Suspicious region, cause known | 106 | rename | A region of a plate with many sample mix-ups, and the cause of the mix-ups could be confidently determined. There was enough information to confidently rename these samples. |
| Cross-cage | 64 | discard | Scattered samples whose predicted mouse is not in the same cage. These mix-ups could not be the result of coprophagy and are therefore very likely to have been caused by a mistake during data generation. |
| Very low uniqueness | 13 | discard | Samples with borderline mix-up results that also looked unusually similar to another sample. The two possible explanations for this situation are 1) the same DNA was added to two wells during library preparation, or 2) well-to-well contamination. |
| Animal swap | 5 | rename | These correspond to the known swap of mice AL-0097 and AL-0105. |
| **Total** | **775** | **discard** |  |
|  | **111** | **rename** |  |

**Supplementary Table 2.** Specific mix-up events.

| Final plate | Action | Type of mix-up | n (n including redoes) | Description |
| --- | --- | --- | --- | --- |
| 7 | rename | Suspicious region, cause known | 14 | Pipetting error during library preparation. Samples were offset by one column. |
| 12 | discard | Suspicious region, cause unknown | 11 (13) | One region of the plate was mostly mix-ups. Cause unknown. **Libraries were unsuccessfully remade for these samples (see final plate 47).** |
| 13 | discard | Suspicious region, cause unknown | 45 (60) | Left half of the plate was mostly mix-ups. Cause unknown. **Libraries were unsuccessfully remade for these samples (see final plate 47).** |
| 15 | discard | Suspicious region, cause unknown | 43 (44) | Bottom half of the plate was mostly mix-ups. Cause unknown. **Libraries were unsuccessfully remade for these samples (see final plate 48).** |
| 16 | discard | Suspicious region, cause unknown | 7 | All samples in row H (except one) were mix-ups. Cause unknown. |
| 18 | discard | Suspicious region, cause unknown | 16 | One region of the plate was mostly mix-ups. Cause unknown. |
| 19 | discard | Suspicious region, cause unknown | 27 | Bottom 3 rows of the plate were mostly mix-ups. Cause unknown. Likely that a mistake happened during EB60. |
| 20 | discard | Suspicious region, cause unknown | 26 | Top 3 rows of the plate were mostly mix-ups. Cause unknown. Likely that a mistake happened during EB60. |
| 25 | rename | Suspicious region, cause known | 36 (37) | Problem with EB74. Some samples were offset by 2 positions, others by 1 position. Renamed contiguous samples with the same offset, kept samples that found the correct mouse, and discarded the rest. |
|  | discard | Suspicious region, cause unknown | 14 (17) |  |
| 27 | rename | Suspicious region, cause known | 7 | Discovered a note in a technician's lab notebook indicating that a row of partially labeled, closed tubes (every other tube was labeled) had been dropped during DNA extraction. Tubes were replaced based on the technician's best guess. Samples were renamed based on the results of mbmix. |
| 28 & 29 | rename | Suspicious region, cause known | 17 | Appears that 4 sets of 6 tubes each were incorrectly rearranged during DNA extraction. Samples were renamed based on the results of mbmix. |
| 31 | discard | Index collision | 77 (136) | Index collision between LB46 (left half of plate) and LB83 (right half of plate). All samples from this plate were discarded, even if the mbmix result was okay. |
|  |  |  |  | LB46 redo samples also collided with other redo samples that had to be discarded. **Libraries were successfully remade for these samples.** |
| 32 | rename | Suspicious region, cause known | 12 | One region from right half of the plate was mostly mix-ups. Most samples were offset by 1, so these were renamed. One sample was offset by -1. This sample was discarded. |
|  | discard | Suspicious region, cause unknown | 1 |  |
| 42 | rename | Suspicious region, cause known | 19 | Bottom 2 rows were all offset in the same way; they contained the mouse in a well one column to the left and five rows up. These samples were renamed. Some samples corresponded to a library well that itself looked like a mix-up, so these samples were discarded. |
|  | discard | Suspicious region, cause unknown | 3 |  |
| 47 | discard | Suspicious region, cause unknown | 49 | Top half of plate and row H were mostly mix-ups. Top half was redo of LB36 (final plate 13), row H was redo of LB32 (final plate 12). |
| 48 | discard | Suspicious region, cause unknown | 42 | Bottom half of plate was mostly mix-ups. Bottom half was redo of LB47 (final plate 15). |

